# TORC1 is an essential regulator of nutrient-dependent differentiation in *Leishmania*

**DOI:** 10.1101/2022.10.20.513059

**Authors:** Elmarie Myburgh, Vincent Geoghegan, Eliza V.C. Alves-Ferreira, Y. Romina Nievas, Jaspreet S. Grewal, Elaine Brown, Karen McLuskey, Jeremy C. Mottram

## Abstract

*Leishmania* parasites undergo differentiation between various proliferating and non-dividing forms to adapt to changing host environments. The mechanisms that link environmental cues with the parasite’s developmental changes remain elusive. Here, we report that *Leishmania* TORC1 is a key environmental sensor for parasite differentiation in the sand fly-stage promastigotes and for replication of mammalian-stage amastigotes. We show that *Leishmania* RPTOR1, interacts with TOR1 and LST8. We investigate TORC1 function by conditional deletion of *RPTOR1*, where under nutrient rich conditions RPTOR1 depletion results in decreased protein synthesis and growth, G1 cell cycle arrest and premature differentiation from proliferative promastigotes to non-dividing mammalian-infective metacyclic forms. These parasites cannot develop into proliferative amastigotes in the mammalian host, or respond to nutrients to differentiate to proliferative retroleptomonads, which are required for their blood-meal induced amplification in sand flies and enhanced mammalian infectivity. RPTOR1-dependent TORC1 functionality represents a critical mechanism for driving parasite growth and proliferation.

## INTRODUCTION

*Leishmania* parasites are responsible for a group of neglected tropical diseases, termed leishmaniases. Their clinical manifestations span a broad range of severity and include self-resolving cutaneous ulcers, debilitating mucocutaneous lesions and lethal systemic disease. The disease group affects the poorest communities with an estimated 700 000 - 1 million new cases each year, and around 1 billion people at risk of infection^1^. The causative protozoan parasites of over 20 *Leishmania* species are transmitted between mammalian hosts by bites of infected female phlebotomine sand flies. The cycling between a mammalian host and insect vector, and movement between different niches within a single host, expose parasites to dramatic changes in environment. In the sand fly, *Leishmania* migrate through different parts of the digestive tract and develop from proliferating procyclic promastigotes to infective non-dividing metacyclic promastigotes that are pre-adapted for survival in their mammalian host^2, 3^. After transmission to the new host these highly motile flagellated promastigotes again experience changes in temperature, pH, nutrients and host-derived factors, and transform to intracellular amastigotes. Successful transmission and survival of parasites relies on their rapid adaptation to these many changes through efficient sensing of the environment and triggering of the appropriate response to drive proliferation, differentiation and/or quiescence.

Eukaryotes, including plants, yeasts, worms, flies and mammals control cell growth and differentiation in response to nutrients and environmental cues through the atypical serine/threonine kinase, TOR (target of rapamycin)^4^. TOR is highly conserved and is found in multiprotein TOR complexes (TORCs) which differ in their components, regulation and function^5^. In mammals, a single TOR forms part of two complexes, TORC1 and TORC2, while the yeast *Saccharomyces cerevisiae* has two TOR orthologues that participate in TORCs with equivalent functions to those in mammals^6^. *Leishmania*, like the closely related kinetoplastid *Trypanosoma brucei*, has four TOR orthologues, TOR1 to TOR4. TOR1 and TOR2 are essential in *Leishmania* promastigotes and have not been characterized, while TOR3 is required for mammalian infectivity and acidocalcisome biogenesis^7, 8^. *Leishmania* TOR4 remains unexplored but it was shown in *T. brucei* to form part of a complex, TORC4, which negatively regulates differentiation to quiescent stumpy form parasites that are pre-adapted for infection of their tsetse fly vector^9^. TORC1 has been studied extensively in many eukaryotic organisms; it senses intracellular nutrient and energy levels to regulate anabolic and catabolic processes. In addition to TOR and its constitutive binding partner, mLST8 (mammalian lethal with Sec13 protein 8), TORC1 contains a complex-specific 150-kDa protein named RPTOR (regulatory-associated protein of mTOR, Kog1 in S. cerevisiae). This adapter protein most likely arose with TOR in the last eukaryotic common ancestor (LECA) and has been conserved with TOR during further evolution of eukaryotes^10, 11^. Due to its role in recruitment of substrates for phosphorylation by TOR, RPTOR is essential for TORC1 functions^12, 13^.

In this study, we defined the TORC1 complex in *Leishmania* and investigated its function through the DiCre-based conditional gene deletion^14, 15^ of *RPTOR* (named *RPTOR1)*. Our proteomic analyses demonstrate that *Leishmania* RPTOR1 interacts with TOR1 and LST8, confirming it as a component of *Leishmania* TORC1. Deletion of *RPTOR1* reveals that TORC1 is required for cell proliferation and long-term survival of promastigotes *in vitro*, and is critical to establishing infections *in vivo*. Notably, *RPTOR1* deletion caused differentiation of procyclic promastigotes to non-dividing metacyclic promastigotes and prevented dedifferentiation of metacyclic promastigotes to dividing retroleptomonads. In addition, complementation with mutant versions of RPTOR1 revealed that a potentially catalytic histidine cysteine dyad in its caspase-like domain is not required for RPTOR1 function showing that it is a pseudopeptidase.

## RESULTS

### *Leishmania* RPTOR1 interacts with TOR1

To define the TORC1 complex in *Leishmania* we generated *L. mexicana* lines expressing N-terminally Twin-Strep-tagged RPTOR1 (LmxM.25.0610), TOR1 (LmxM.36.6320) or the control bait LmxM.29.3580 using a CRISPR-Cas9 mediated endogenous tagging approach^16^. RPTOR1 or TOR1 were enriched from parasite lysates with Streptactin-XT, allowing specific elution with biotin and quantitative analysis of complexes by mass spectrometry (Figure 1A, Figure S1A and Table S1). A RPTOR1-TOR1 interaction was detected in both pulldowns and LST8 co-enriched with both RPTOR1 and TOR1 indicating that *Leishmania* RPTOR1 associates with two proteins that are key components of TORC1 in other eukaryotes. The TOR1 pull-down also detected an HSP90 chaperone (referred to as HSP83-1), the co-chaperone HIP (HSC-70 interacting protein, LmxM.08_29.0320) and two additional proteins that have been linked to TOR signalling or regulation in other organisms, RUVBL1 (LmxM.33.3500) and RUVBL2 (LmxM.33.2610), which are homologues of the ATPases Pontin (RUVBL1) and Reptin (RUVBL2). Interestingly, two proteins with no known function (LmxM.16.1080 and LmxM.08_29.1470) were significantly enriched in both the RPTOR1 and TOR1 purifications, indicating that they might form part of the core TORC1 complex in *Leishmania*. Homology and protein domain searches with the 646 amino acid sequence of LmxM.16.1080 revealed homologous sequences in *Leishmania* and *Leptomonas spp*; however, no protein domains were predicted. LmxM.08_29.1470 contains an ankyrin repeat domain suggesting a role in protein-protein interaction, and orthologues of this 711 amino acid protein are present in several kinetoplastids including *Trypanosoma*, *Leishmaniiae*, *Strigamonadinae and Bodo saltans*.

**Figure 1.**
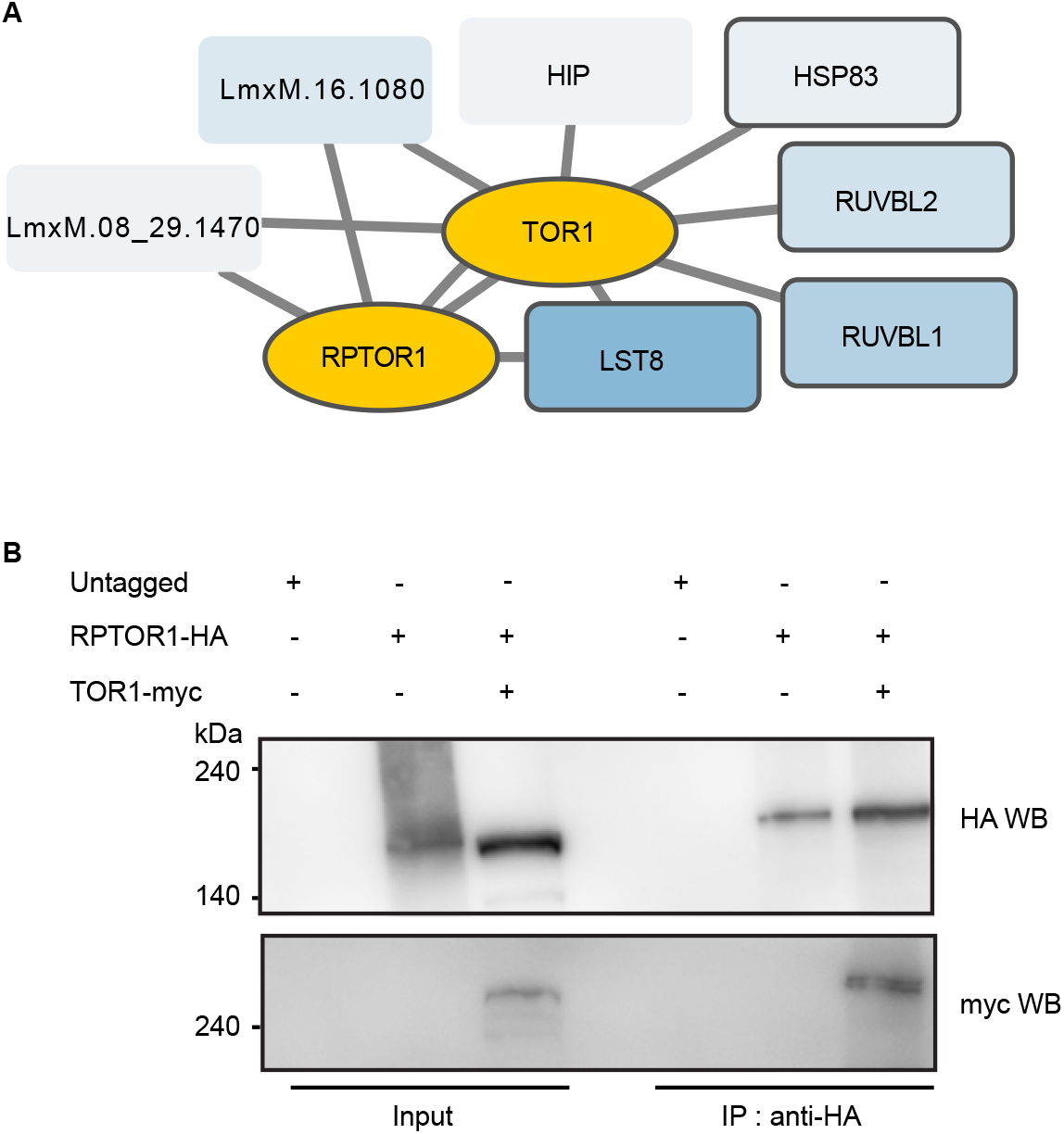
Characterization of the RPTOR1 containing complex in *Leishmania*. (*A*) Proteins associated with RPTOR1 and TOR1 were identified by mass spectrometry analyses of strep-tagged RPTOR1 or TOR1 immunoprecipitates. Non-specific interactors were removed using stringent filtering criteria (see Methods) and SAINT analyses. Bait protein are shown in yellow and identified interactors are in blue. Solid outline show proteins previously identified as interactors in mass spectrometry with human proteins. (*B*) Validation of RPTOR1-TOR1 interaction by co-immunoprecipitation. Lysates of *L. mexicana* cells expressing untagged RPTOR1 and TOR1, HA-tagged RPTOR1 and /or myc-tagged TOR1 were incubated with anti-HA- or anti-myc-conjugated magnetic beads and analysed by western blot (WB) using anti-myc or anti-HA antibodies.

To confirm the interaction of RPTOR1 and TOR1 we co-expressed RPTOR1-HA and TOR1-Myc in *L. mexicana* and performed western blotting on immuno-precipitated complexes. *L. mexicana* with untagged RPTOR1, or parasite lines singly expressing RPTOR1-HA (Figure 1B) or TOR1-Myc (Figure S1B) were included as controls. Our analysis showed that TOR1-Myc is immunoprecipitated with RPTOR1-HA confirming that these two proteins are part of a TORC1 complex.

### RPTOR1 is essential for cell proliferation

Previous unsuccessful attempts to generate a *TOR1* null mutant in *L. major* and *L. mexicana* suggests that TOR1 is essential for *Leishmania* promastigotes^7, 8^. To investigate whether RPTOR1 is also essential for *Leishmania* promastigotes, we attempted to generate *RPTOR1* null mutants in *L. major* and *L. mexicana*. Multiple attempts to replace *RPTOR1* in *L. major* using standard homologous recombination and in *L. mexicana* using CRISPR-Cas9 mediated homologous recombination failed to generate homozygous knockout mutants despite the generation of heterozygous mutants. These observations suggested indirectly that RPTOR1 may be essential for promastigote survival. To provide direct evidence for essentiality and explore the functional consequence of *RPTOR1* deletion we made use of a DiCre conditional gene deletion system^14, 15^ which excises *LoxP* flanked (floxed) *RPTOR1* following rapamycin treatment (Figure S1B). A DiCre expressing *L. major* cell line (DiCre) was modified to replace one *RPTOR1* allele with a floxed GFP-tagged *RPTOR1* expression cassette (*RPTOR*^+/flox^) and the 2^nd^ allele with an antibiotic (hygromycin) resistance cassette (*RPTOR1*^−/flox^). Diagnostic PCR amplification confirmed the integration of the knockout and expression cassettes in several hygromycin resistant clones (Figure S1C). Two of these clones, clones 1 and 2, were selected for further analysis, and PCR showed that rapamycin treatment of these *RPTOR1*^−/flox^ promastigotes for 3 days induced excision of the *RPTOR1* coding sequence (CDS) (Figure 2A and B). Rapamycin is well known as an inhibitor of mammalian TOR, however in trypanosomatids the residues important for rapamycin binding are mutated, making them relatively insensitive to rapamycin inhibition of TOR^7, 17^. To ensure we could distinguish between RPTOR1 deletion and other potential rapamycin-specific effects, we also included the control cell line (DiCre) treated with rapamycin in all experiments. Flow cytometry of live promastigotes indicated a complete loss of RPTOR1-GFP protein in inducible knockout cell lines (*RPTOR1*^−/flox^ clones 1 and 2) following rapamycin treatment (Figure 2C and D). The GFP signal in both clones was reduced to the level detected in the DiCre control cells.

**Figure 2.**
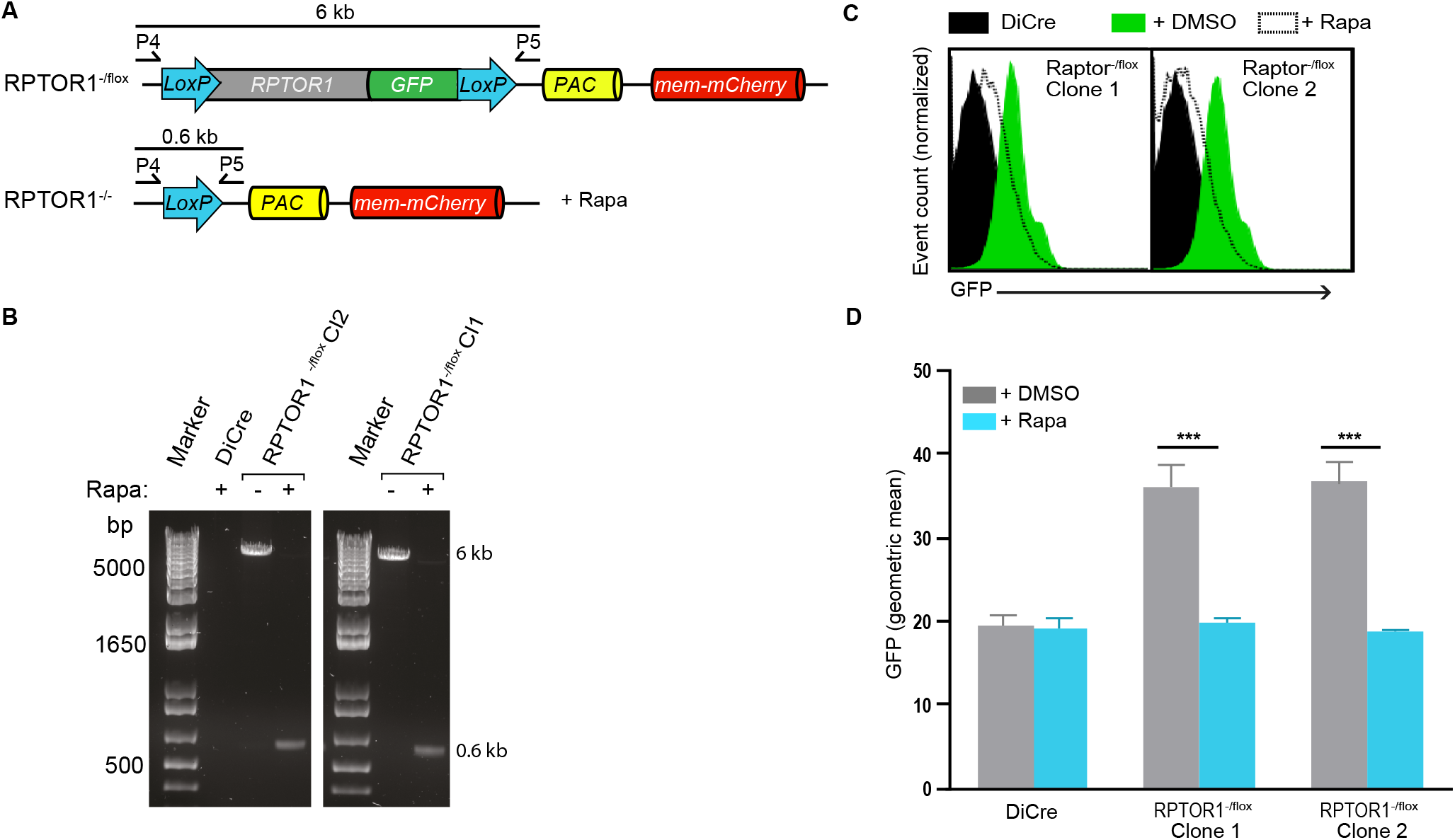
Generation of inducible *RPTOR1* knockout lines. (*A*) Schematic of the *RPTOR1^−/flox^* locus and the expected *RPTOR1^−/−^* locus following rapamycin-induced recombination. Oligonucleotide binding sites and the length of the PCR amplicons to detect *RPTOR1* excision are indicated. (*B*) PCR analysis of genomic DNA from DiCre and inducible *RPTOR1^−/flox^* clonal lines (Cl1 and Cl2) showing recombination after three days of treatment with 100 nM Rapamycin (Rapa) added daily. (*C*) Analyses of GFP-tagged RPTOR1 loss by flow cytometry. Representative histograms show GFP fluorescence in live *L. major* promastigotes from untreated (+DMSO, green with solid line) and rapamycin (+Rapa, white fill with dotted line)-treated cell lines. Rapamycin-treated DiCre (black fill) is included in each histogram as GFP negative control. Cells were treated with DMSO or rapamycin for three days, diluted and cultured for another three days before analysis by flow cytometry. (*D*) The GFP geometric mean fluorescence intensity in untreated (+DMSO) or rapamycin (+Rapa)-treated cells shown in C. Data represent mean± SEM from three individual experiments; \**P* value ≤ 0.05, \*\*\**P* value ≤ 0.001 in an unpaired t test.

To characterise the essential role of RPTOR1 for *Leishmania* survival we started our analyses by looking at parasite proliferation. Excision of *RPTOR1* resulted in a significant decrease in promastigote proliferation compared to uninduced (DMSO-treated) *RPTOR1*^−/flox^ cells and the parental cell line, DiCre (Figure 3A and Figure S2A). Cell densities in the rapamycin-treated lines were 30–40% lower on day 4 and 70–80% lower by day 7 after treatment compared to the controls. To assess the effect of RPTOR1 loss on long-term parasite survival we used a clonogenic assay and found that rapamycin-induced RPTOR1 depletion caused an 80–96% reduction in clone survival (Figure 3B). Furthermore, PCR analysis showed that all surviving clones retained a copy of *RPTOR1* (Figure S2B) confirming an essential role for RPTOR1 in parasite survival. We next explored the reason for reduced cell proliferation and survival in the absence of RPTOR1. Flow cytometry of live cells stained with propidium iodide indicated that *RPTOR1* excision did not cause a significant amount of cell death on days 5 and 6 after induction (Figure 3C and Figure S3A). However, induced *RPTOR1*^−/flox^ cells arrested in the G1 phase of the cell cycle; there was a 26–32 % increase of cells in G1, a 39–46 % reduction in S and a 19–32 % reduction in G2/M compared to the uninduced control cells on day 5 (Figure 3D). At later time points (day 8) both uninduced and rapamycin-induced DiCre and *RPTOR1*^−/flox^ lines were arrested in G1, consistent with the predicted development of non-proliferative metacyclic promastigotes in a late stage nutrient-depleted *in vitro* culture (Figure S3B)^18^. Collectively, these results suggest that RPTOR1 plays an essential role in promastigote cell proliferation and survival, and that its loss results in cell cycle arrest in G1. Importantly, we observed the same cell proliferation defect and cell cycle arrest in our two independently generated *RPTOR1*^−/flox^ clones.

**Figure 3.**
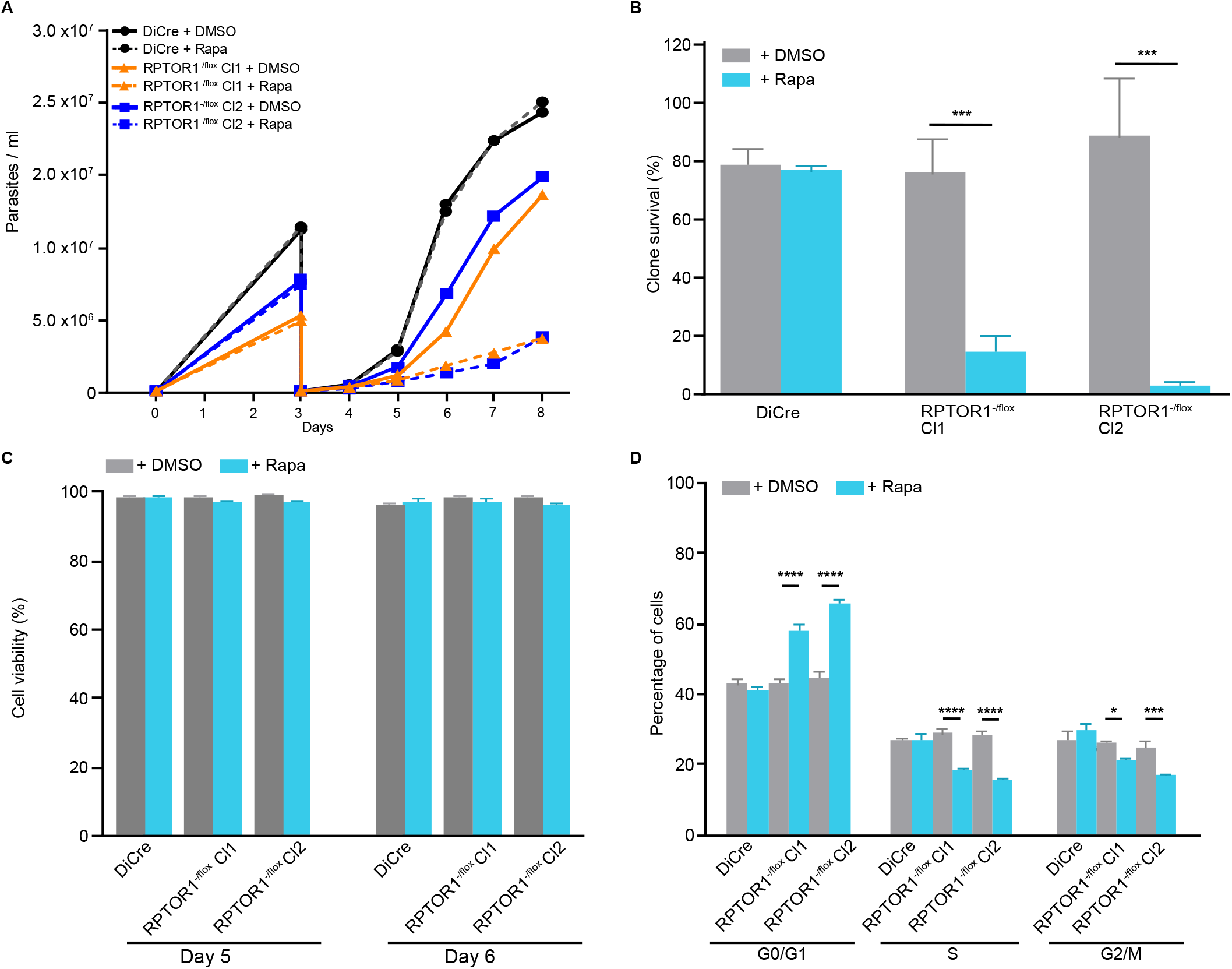
RPTOR1 is essential for cell proliferation. (*A*) Cell densities of rapamycin-induced relative to uninduced cells (shown as ratio to uninduced). Log-stage promastigotes of DiCre and *RPTOR1^−/flox^* lines (Cl1 and Cl2) were set up at 1×10^5^ cells mL^−1^ (day 0) and treated for three days with 0.1% DMSO or 100 nM rapamycin; cells were then counted and diluted to 1×10^5^ cells mL^−1^ (day 3) followed by culturing and daily counting for five days. Data represent mean± SD from three to four individual experiments. \**P* value ≤ 0.05, \*\*\**P* value ≤ 0.001 in a one-way ANOVA with Tukey’s post hoc test. (*B*) Clone survival (shown as a percentage of plated clones) of uninduced (+DMSO) and rapamycin-induced (+ Rapa) cells. Data represent mean± SD from three to four individual experiments with two to four technical replicates each. \*\*\**P* value ≤ 0.001 in a one-way ANOVA with Tukey’s post hoc test. (*C*) Cell viability of uninduced and rapamycin-induced cells. Log-stage promastigotes were treated for 3 days, then diluted and cultured for another two or three days with DMSO or rapamycin. Cells were stained with propidium iodide and analysed by flow cytometry. (*D*) Cell cycle analysis of uninduced and rapamycin-induced cells from DiCre, *RPTOR1^−/flox^* Cl1 and *RPTOR1^−/flox^* Cl2 lines. Promastigotes were fixed, stained with propidium iodide and analysed by flow cytometry. Data represent mean± SD of replicate samples from a representative experiment. \**P* value ≤ 0.05, \*\*\**P* value ≤ 0.001, \*\*\*\**P* value ≤ 0.0001 in a two-way ANOVA with Bonferroni post hoc test.

### RPTOR1 controls protein synthesis and cell size

TORC1 signalling in humans regulates protein synthesis and cell growth. We asked whether RPTOR1 depletion impacts these essential processes in *Leishmania*. Protein synthesis, measured by incorporation of the synthetic methionine analogue L-azidohomoalanine (AHA), was reduced by 51% and 90% in *RPTOR1*^−/flox^ clones 1 and 2, respectively following rapamycin-induced loss of RPTOR1 (Figure 4A). To analyse cell growth, the size of promastigote parasites was measured by forward light scatter using flow cytometry on day 5 and 8 after induction. In this protocol cells from log-phase promastigote cultures were diluted in fresh nutrient rich medium on day 3 so that cells should be actively proliferating and exhibiting normal growth at day 5. By day 8 media is more depleted of nutrients and parasites would have differentiated to metacyclic promastigotes which are smaller in size compared to the proliferating procyclic promastigotes of day 5. Forward scatter was reduced following *RPTOR1* excision (Figure 4B and C); on day 5 the geometric means in *RPTOR1*^−/flox^ lines decreased by 60–65 % compared to the uninduced controls, and by 37–43% compared to the DiCre control. Two cell populations based on size were visible in the induced *RPTOR1*^−/flox^ lines with 23.6–40.4 % and 71.4–78.7% smaller cells measured on day 5 and 8, respectively. By day 8 the DiCre cells had also reduced in size consistent with the development of smaller metacyclic promastigotes, but these cells were still larger than the cells measured in the *RPTOR1*^−/−^ lines. Interestingly, the uninduced *RPTOR1*^−/flox^ lines had increased forward scatter compared to the DiCre line at both day 5 and day 8 (Figure 4C). We next measured cell size by scanning electron microscopy (SEM) of one of our *RPTOR1*^−/flox^ clones (clone 2) (Figure 4D). SEM images revealed that the difference in cell size observed by flow cytometry is related to a decrease in body width of parasites (mean ± SD, 1.5 ± 0.3 μm in rapamycin-induced compared to 2.0 ± 0.3 μm in uninduced). Additionally, body and flagellum length were increased in *RPTOR1*^−/−^ cells with body lengths of 9.4 ± 2.8 μm for rapamycin-induced compared to 7.5 ± 1.9 μm for uninduced cells and flagellum lengths of 13.8 ± 4.2 μm for rapamycin-induced compared to 8.1 ± 3.9 μm for uninduced cells (Figure 4D and E). Rapamycin treatment of DiCre control cells resulted in an increase in body width and a decrease in flagellum length. These results indicate that RPTOR1 is important for promastigote protein synthesis and normal cell growth.

**Figure 4.**
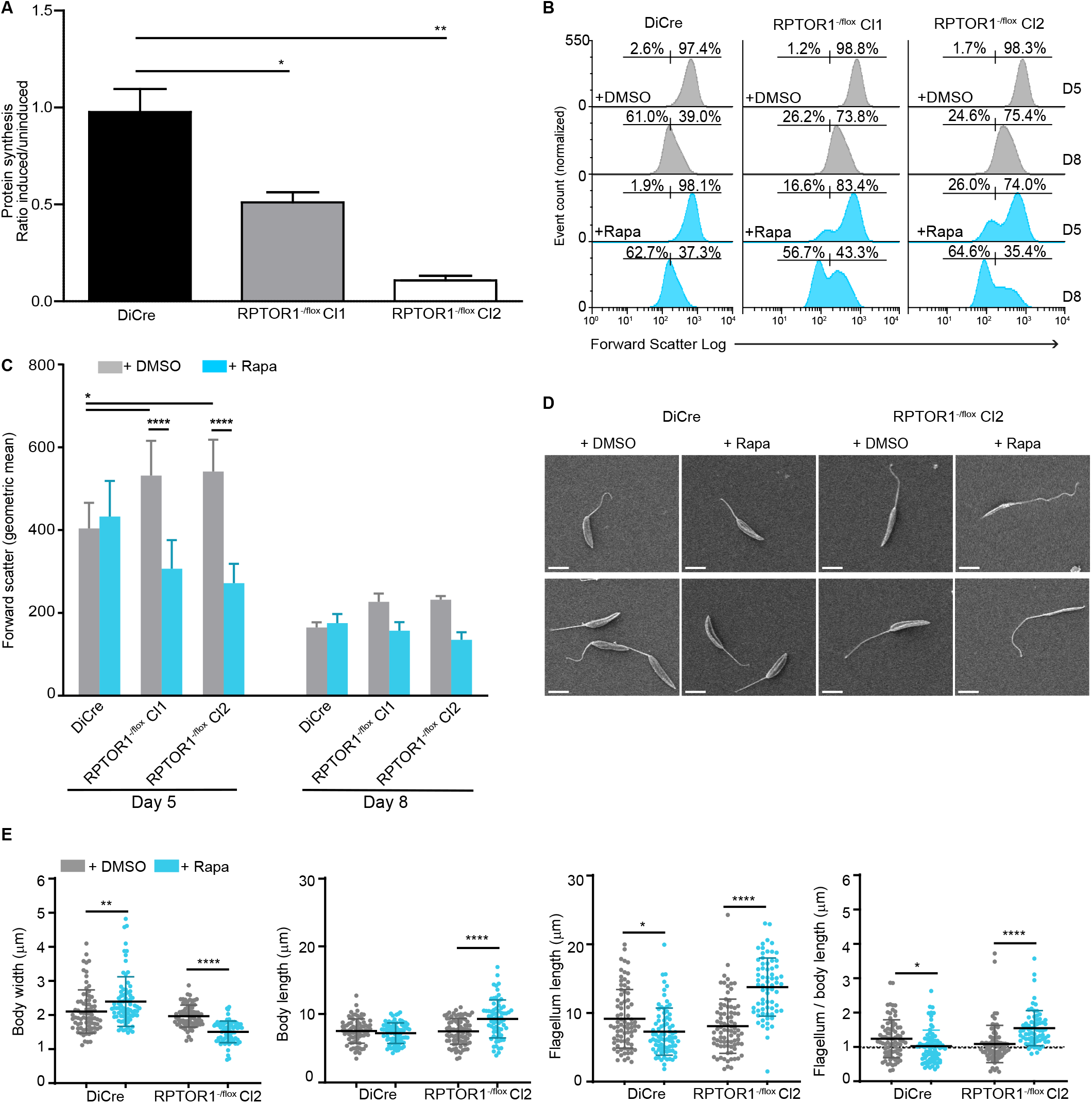
RPTOR1 controls protein synthesis and cell size. (*A*) Reduction in protein synthesis following rapamycin-induced *RPTOR1* excision. Cells were harvested after six days of daily induction. Newly synthesised proteins were measured by AHA incorporation using a CLICK-It assay. The ratio of AHA incorporation in rapamycin-induced relative to uninduced (DMSO-treated) cells is shown. Data represent mean± SEM from four individual experiments. (*B*) Cell size was measured by flow cytometry in fixed cells after five (D5) or eight (D8) days of treatment with DMSO or rapamycin (Rapa). Histograms of forward scatter from one representative sample of each condition are shown to indicate cell size. Percentages of smaller versus larger cells are shown above the gates for each of these populations. (*C*) Forward scatter fluorescence intensity of indicated lines after five and eight days of treatment. Data represent geometric mean ± SD of pooled data from 4 individual experiments. (*D*) Scanning electron microscopy images of DiCre and *RPTOR1^−/flox^* Cl2 promastigotes after five days of treatment with DMSO or rapamycin. Scale bar = 5 μm (*E*) Body width, body length and flagellum length were measured in 70-80 individual cells. Means ± SD are shown, and each dot represents the measurement in a single cell. For *A to E* cells were diluted in fresh medium after the first three days of treatment. \**P* value ≤ 0.05, \*\**P* value ≤ 0.01, \*\*\**P* value ≤ 0.001, \*\*\*\**P* value ≤ 0.0001 in a one-way ANOVA with Tukey’s post hoc test.

### Functional analysis of RPTOR1 caspase domain

RPTOR1 is a large protein molecule consisting of 1471 residues, composed of the characteristic domains of human RPTOR1 (Figure 5A). It features a raptor N-terminal conserved (RNC) region (Raptor N-terminal Caspase-like domain), with ~38 % identity to the human RNC, an Armadillo repeat domain and C-terminal WD40 repeats, both implicated in protein-protein interactions. The RNC region is structurally similar to caspases^19^; it contains an α-β-α sandwich fold with an apparent histidine-cysteine (His-Cys) dyad near the caspase active site. As there are no three-dimensional structures of LmRPTOR1, a structural alignment and topological comparison (Figure 5B) of *Arabidopsis thaliana* RAPTOR1 RNC (AtRAPTOR1) with human Caspase-7 (*Hs* Caspase-7) was carried out. This showed that the secondary structural elements of AtRAPTOR1 and HsCaspase-7 are highly conserved, along with the position of the catalytic histidine and the glycine that follows it. However, in AtRAPTOR1 a serine (Ser^246^) directly aligns with the Caspase-7 active site cysteine although a cysteine (Cys^245^) does directly precede it (Figs. 5B and C and Figure S4A)^20^. Primary sequence alignment of RPTOR1 from *A. thaliana*, *H. sapiens* and *L. major* indicate that these regions are also conserved in the human and *L. major* proteins (Figure 5C and Figure S4B) suggesting that the cysteine of the His-Cys dyad is also shifted in these other RPTOR1 proteins, with an asparagine residue (Asn^270^) occupying the position of the catalytic cysteine in *Lm* RPTOR1 (Figure 5B). Structural superposition of the Alphafold model of *L. infantum* RPTOR1 (LINF_250011400), AtRAPTOR1 and HsCaspase-7 using UCSF Chimera confirmed our results above (Figure S5).

**Figure 5.**
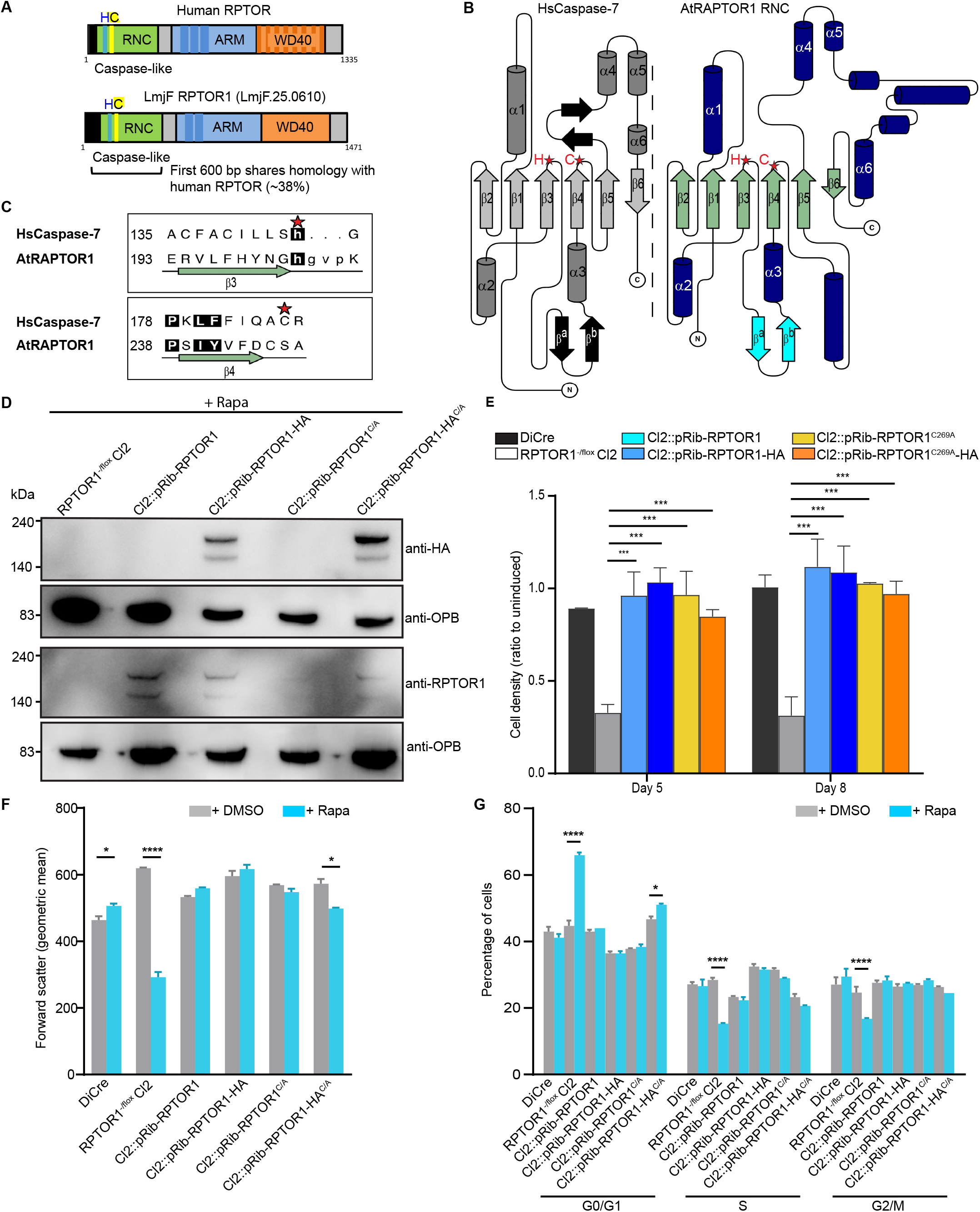
Functional analysis of RPTOR1 caspase domain. (*A*) Schematic representation of human RPTOR and *L. major* RPTOR1 showing the caspase-like raptor N-terminal conserved (RNC) domain, armadillo repeat (ARM) and C-terminal WD40 repeat domains. The potential caspase active site histidine (H) cysteine (C) dyad is highlighted. (*B*) Topology diagrams showing structural similarities between the caspase-like domain of human caspase-7 (left) and *A. thaliana* RPTOR1 (right). β-strands are coloured light grey or green, and α-helices are coloured dark grey or dark blue. Conserved structural elements are numbered from the N terminus and the and the position of the histidine-cysteine dyad is shown by red stars. (*C*) Sequence alignment of the histidine-cysteine dyad containing regions from human caspase-7 and *A. thaliana* RAPTOR1. β-strands are coloured green; loop regions shown as a black line and the histidine and cysteine residues of the dyad are indicated for caspase-7 by red stars. Residues where the backbones of the two structures overlay tightly are shown in upper case and those with higher deviation/no matching residue are in lower case. Numbers at the start of each line indicate the position of the first residue shown in that line in the UniProt sequences. (*D*) Western blot analysis of lysates from rapamycin-induced (+ Rapa) cell lines confirms the expression of untagged and HA-tagged RPTOR1 in the complementation mutants and RPTOR1’s absence in the knockout mutants. Cells were induced for 3 days, diluted in fresh culture media and induced for another 3 days. (*E*) Cell densities of rapamycin-induced relative to uninduced DiCre, *RPTOR1^− /flox^* Cl2 and *RPTOR1* complementation lines (shown as ratio to uninduced) five days or eight days after induction. The graph shows the mean± SD of pooled data from two individual experiments. \*\*\**P* value ≤ 0.001 in a two-way ANOVA with Bonferroni post hoc test. (*F*) Forward scatter fluorescence intensity of indicated lines after five days of induction. Data represent geometric mean± SD of replicate samples from one representative experiment. (*G*) Cell cycle analysis of uninduced and rapamycin-induced cells. Promastigotes were fixed, stained with propidium iodide and analysed by flow cytometry. Data represent mean± SD of replicate samples from one representative experiment of two. \**P* value ≤ 0.05, \*\*\**P* value ≤ 0.001 in a two-way ANOVA with Bonferroni post hoc test.

To assess whether *Leishmania* RPTOR1 requires protease activity from the caspase-like RNC for its function we generated addback lines in *RPTOR1*^−/flox^ clone 2 by integration of untagged and HA-tagged wildtype and active site mutant *RPTOR1* in the ribosomal locus. For the active site mutant, the cysteine at position 269 in the predicted His-Cys dyad was mutated to an alanine (C269A). This resulted in the creation of Cl2*∷pRib-RPTOR1*, Cl2*∷pRib-RPTOR1-* HA, Cl2*∷pRib-RPTOR1^C269A^* and Cl2*∷pRib-RPTOR1-HA^C269A^*. Western blot analyses were performed with either anti-HA antibodies or antibodies raised against a recombinant 462 amino acid *L. major* RPTOR1 fragment. This detected expression of RPTOR1 or RPTOR1-HA following rapamycin-induced excision of floxed *RPTOR1* in the endogenous locus of the addback lines (Figure 5D). Expression of RPTOR1, RPTOR1-HA, RPTOR1^C269A^ or RPTOR1-HA^C269A^ restored parasite proliferation following rapamycin-induced excision of floxed *RPTOR1* (Figure 5E). We also observed rescue of the cell size defect and G1 cell cycle arrest with RPTOR1 or RPTOR1^C269A^ re-expression (Figs. 5F and G). These results indicate that re-expression of RPTOR1 restores the phenotypes observed after rapamycin-induced excision of floxed RPTOR1 and provides confidence that these phenotypes are specific for RPTOR1 deletion. Furthermore, the disruption of the potential catalytic site cysteine in the caspase-like domain does not impact on complementation of RPTOR1 function suggesting that RPTOR1 caspase activity is therefore not required for promastigote proliferation and growth.

### RPTOR1 loss induces parasite differentiation to metacyclic promastigotes

*Leishmania* promastigotes undergo several biochemical and morphological changes in their sand fly hosts to eventually differentiate into mammalian-infective metacyclic promastigotes. This differentiation process, termed metacyclogenesis, is induced *in vitro* by low pH, depletion of adenosine and low levels of tetrahydrobiopterin^21, 22, 23, 24^ but the mechanisms involved and the exact triggers for differentiation remain poorly defined. Our data showed that loss of RPTOR1 resulted in an increase in parasites with characteristics of metacyclic promastigotes: cells were non-proliferative, arrested in G1 and had relatively long flagella. Manual counting of live cells also revealed that *RPTOR1*^−/−^ parasites were highly motile. Since RPTOR1 has been shown to play a role in nutrient sensing in other organisms, we investigated whether it may be involved in nutrient sensing to coordinate continued proliferation or metacyclogenesis in *Leishmania*. We assessed whether *RPTOR1* deletion triggered metacyclogenesis by measuring modification of lipophosphoglycan (LPG) and expression of metacyclic stage-specific proteins. LPG structure is modified during metacyclogenesis leading to a decrease in agglutination of parasites by peanut lectin (PNA)^25^. As a positive control we included stationary-phase DiCre cells that had been grown for seven days to indicate the percentage of metacyclic promastigotes purified from a nutrient-depleted culture for each assay. Rapamycin-induced excision of *RPTOR1* in *RPTOR1^−/flox^* resulted in a 14–18-fold increase in non-agglutinated (PNA^−^) parasites compared to DiCre after three days of culture in nutrient-rich medium, with 3.5% and 2.7% PNA^−^ cells in *RPTOR1^−/flox^* clone 1 and 2, respectively (Figure 6A). In contrast, rapamycin-induced DiCre grown in the same nutrient-rich conditions had only 0.19% PNA^−^ cells compared to 0.33% in uninduced DiCre cells. Re-expression of RPTOR1 or RPTOR1^C269A^ restored the phenotype back to levels in the DMSO-treated *RPTOR1*^−/flox^ cells, confirming once again that this potential catalytic site is not required for RPTOR1’s role in cell proliferation. Furthermore, *RPTOR1*^−/−^ cells that had been continuously cultured for seven days with rapamycin had 2.6- and 4-fold more PNA^−^ cells in clone 1 and 2, respectively, compared to DiCre stationary-phase cells (also grown for seven days) (Figure 6B). Interestingly, DMSO-treated *RPTOR1*^−/flox^ cells had significantly less PNA^−^ cells than DMSO-treated DiCre for both the 3 and 7-day culture conditions, with ~0.002% and 0.2– 1.4% PNA^−^ for *RPTOR1*^−/flox^ cells in the respective culture conditions compared with 0.3% and 8.4% for DiCre cells (Figure 6A and B). This suggests that *RPTOR1*^−/flox^ cells show a reduction in metacyclogenesis compared to control DiCre cells. We next measured expression of the metacyclic stage-specific protein, SHERP (small hydrophilic endoplasmic reticulum-associated protein)^26^ using flow cytometry. Loss of RPTOR1 resulted in a significant increase in cells with high SHERP expression while *RPTOR1*^−/flox^ cells had lower SHERP expression compared to the DiCre control (Figure 6C and Figure S6A and B). These data indicate that loss of RPTOR1 gives rise to parasites with surface characteristics of metacyclic promastigotes. It also suggests that the altered *RPTOR1* gene in the *RPTOR*^−/flox^ line results in inhibition of metacyclogenesis compared to control DiCre cells with a WT *RPTOR1* gene.

**Figure 6.**
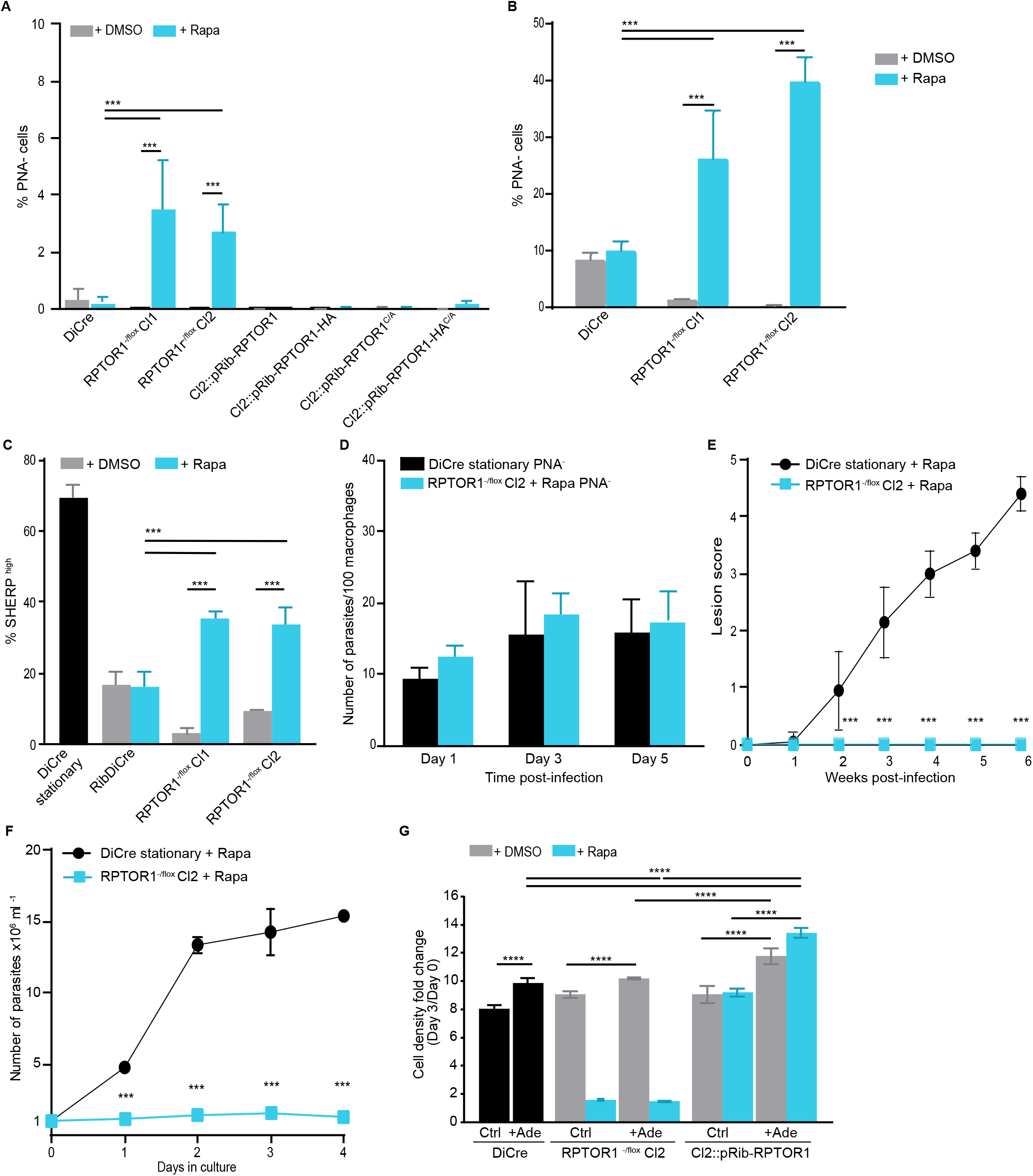
RPTOR1 loss induces metacyclogenesis but is detrimental for murine infection. (*A*) Percentage of metacyclic promastigotes (PNA^−^ cells) in cultures treated daily for six days with DMSO or 100 nM rapamycin (Rapa). Cells were diluted in fresh medium after the first three days of treatment. RPTOR1 loss increases the percentage of PNA^−^ cells while complementation rescues the phenotype. (*B*) Percentage of metacyclic promastigotes in cells cultured continuously for seven days with daily addition of DMSO or rapamycin (Rapa). The graphs in *A* and *B* show the mean± SD of pooled data from three to four individual experiments. \*\*\**P* value ≤ 0.001 in a one-way ANOVA with Tukey’s post hoc test. (*C*) *RPTOR1* excision increases the percentage of cells expressing high levels of SHERP. Expression was analysed by flow cytometry in fixed cells stained with anti-SHERP and Alexa Fluor 647-conjugated secondary antibodies. Graph shows mean ± SD from pooled data of three experiments (*D*) Macrophage infectivity of PNA^−^ promastigotes. Bone-marrow macrophages were infected with parasites at a 1:1 ratio and analysed at day 1, 3 and 5 after infection using microscopy. Graph shows mean ± SEM of triplicate wells from one representative of two experiments. (*E*) Lesion development (depicted by lesion score) in BALB/c mice injected intradermally in the ear with 1×10^5^ metacyclic promastigotes (n = 10 mice per group combined from two experiments, mean± SD is shown). \*\*\**P* value ≤ 0.001 in a two-way ANOVA with Bonferroni post hoc test. (*F*) *RPTOR1^−/−^* metacyclic promastigotes are unable to differentiate to proliferating retroleptomonads in nutrient-rich medium. Purified metacyclic promastigotes were cultured in nutrient-rich medium. Data represent mean± SD of triplicate cultures from one of two similar experiments. \*\*\**P* value ≤ 0.001 in a two-way ANOVA with Bonferroni post hoc test. For *A*, *C, D, E and F*, metacyclic promastigotes were purified by PNA agglutination from stationary-phase DiCre cells cultured for 7 days in the presence of rapamycin, and *RPTOR1^−/flox^* cells induced with rapamycin for three days, diluted and cultured for three more days with daily addition of rapamycin. (*G*) Cell proliferation of rapamycin-induced cells from DiCre, *RPTOR1^−/flox^* Cl2 and *RPTOR1* complementation lines grown in Grace’s medium with the addition or not of 500 μM adenine. After rapamycin treatment for five days cells were diluted (day 0) and cultured for a further three days (day 3) in the presence or not of adenine. Proliferation fold change (density at day 3 compared to day 0) is reported. Graph shows mean ± SD of triplicate cultures from one experiment. \*\*\*\**P* value ≤ 0.0001 in a one-way ANOVA with Tukey’s post hoc test.

### Loss of RPTOR1 abrogates lesion development in mice

Next, we investigated the infectivity of these metacyclic promastigote-like cells by purifying PNA^−^ cells from DiCre control or *RPTOR1^−/flox^* after rapamycin induction and assessing infection of bone-marrow derived macrophages. We observed similar infectivity between the DiCre control or *RPTOR1^−/flow^* indicating that *RPTOR1^−/−^* PNA^−^ cells are as infective as control PNA^−^ metacyclic promastigotes (Figure 6D). The time scale of *in vitro* macrophage infections suggests that *RPTOR1*^−/−^ parasites infect macrophages and may differentiate to amastigotes but it doesn’t provide evidence that these parasites can replicate as amastigotes. To address this question, we infected BALB/c mice in the ears with PNA^−^ DiCre or PNA^−^ *RPTOR1*^−/−^ parasites and monitored lesion development. Loss of RPTOR1 completely abrogated lesion development in mice, unlike the DiCre control, showing that despite having the molecular features of metacyclic promastigotes, the *RPTOR1*^−/−^ parasites cannot establish an infection or replicate as amastigotes *in vivo* (Figure 6E).

### RPTOR1 is essential for differentiation of *Leishmania* metacyclic promastigotes to retroleptomonads

Non-dividing *Leishmania* metacyclic promastigotes have been reported to have the ability to differentiate to replicative retroleptomonads in response to a blood meal in the sand fly vector or addition of nutrients in culture^18, 27^. We hypothesized that if RPTOR1 is involved in nutrient sensing to promote cell proliferation then it may also influence this process. Our results showed that while PNA^−^ metacyclic DiCre proliferated in nutrient-rich medium, PNA^−^ *RPTOR1*^−/−^ parasites remained non-proliferative (Figure 6F). These data suggest that RPTOR1 is not only essential for proliferation of leptomonad promastigotes and amastigotes but also for the nutrient-induced differentiation of metacyclic promastigotes to retroleptomonads, a critical process for maintaining infectivity of the insect vector and successful *Leishmania* transmission.

### TORC1 is downstream of the purine sensing mechanism

*Leishmania* cannot synthesise purine nucleotides *de novo* and are dependent on this nutrient’s availability in their host for growth^28, 29^. Purine starvation, for example through the withdrawal of purines *in vitro* from culture medium, results in parasite growth arrest in G1/G0 of the cell cycle and metabolic changes to a quiescent-like state^29, 30^. Conversely, purine supplementation can reverse this growth arrest. Purine sensing has also been linked to metacyclogenesis with adenosine supplementation recovering proliferation of metacyclic promastigotes in culture and inhibiting metacyclogenesis in the sand fly host^24^. We predicted that TORC1 would function downstream of the parasite purine sensing mechanism and may be a key complex to facilitate nutrient sensing, including for purines. To investigate this, we assessed how RPTOR1 deletion influenced parasite growth following adenine supplementation. Addition of adenine increased the proliferation of rapamycin-treated DiCre and uninduced (DMSO-treated) *RPTOR1*^−/flox^ parasites (Figure 6G). On the other hand, *RPTOR1^−/−^* parasites showed a low level of proliferation that did not increase in response to adenine supplementation. We also included our wildtype RPTOR1 re-expression line (Cl2*∷pRib-RPTOR1*) to assess whether higher levels of RPTOR1 in parasites might increase responsiveness to adenine supplementation. In this case, adenine supplementation increased proliferation even more compared to the DiCre and *RPTOR1^−/−^* parasites. This suggests that the level of RPTOR1, and thus TORC1 activity in cells may fine-tune the responsiveness to nutrients with higher levels of RPTOR1 resulting in more proliferation of cells when nutrients are available.

## DISCUSSION

The TORC1 signalling pathway in *Leishmania* is largely unexplored due to its critical role in cell growth and proliferation, and the technical challenges of functionally characterising essential genes in this organism^7, 31^. In this study, we confirm TORC1 essentiality through analysis of the TOR1 complex binding partner, RPTOR1, and use a conditional gene deletion system to reveal an important role for RPTOR1 in the regulation of cell proliferation, differentiation and thus infectivity. A previous report suggested that TORC1 is essential for *Leishmania* promastigote survival due to the inability to generate TOR1 homozygous knockouts^7^. Our results confirmed this essential role of TORC1 for promastigote growth and show that it is also critical for the growth and infectivity of amastigotes *in vivo* and the dedifferentiation of metacyclic promastigotes to proliferative retroleptomonads.

*Leishmania* TORC1 composition, function and regulation is unknown. Bioinformatic analysis by us and others^7, 11, 32^ identified *Leishmania* homologues for the key TOR1 complex members TOR, RPTOR1 and LST8 (also found in TORC2) but not for DEPTOR and PRAS40, which associate with these TORC1 members in vertebrates^4, 33^. In *T. brucei* two distinct TOR1 and TOR2 complexes were identified with TORC1 containing TOR1 and RPTOR, and TORC2 containing TOR2 and RICTOR. Our proteomic analysis of the RPTOR1 and TOR1-containing complexes showed that *Leishmania* RPTOR1 directly associates with TOR1 and LST8 to form TORC1. The RPTOR1-TOR1 interaction was furthermore confirmed by co-immunoprecipitation. Interestingly, our mass spectrometry also identified two proteins of unknown function (Gene IDs: LmxM.16.1080 and LmxM.08_29.1470) that interact with both TOR1 and RPTOR1. Bioinformatic analysis did not reveal any known homologues or published data to predict potential functions for these proteins, so further work is required to establish their role in TOR signalling and parasite growth. Other TOR1 interactors included the AAA+ family ATPases RUVBL1 and RUVBL2, the molecular chaperone HSP83 (an HSP90 homologue) and the co-chaperone HIP, which associates with intermediate HSP90 and HSP70 complexes^34^. In humans RUVBL1 and RUVBL2 form part of several multiprotein complexes that are involved in chromatin remodelling, transcription, telomerase assembly, snoRNP (small nucleolar ribonucleoprotein) biogenesis and phosphoinositide three-kinase-related kinase (PIKK) regulation^35, 36, 37^. Abrogation of either RUVBL protein impairs cell growth and proliferation in several organisms^37^. The *Leishmania* proteins contain the characteristic DNA-binding and ATPase motifs of their human homologues and are predicted to have similar enzymatic activities to these proteins in other organisms^38^ but their importance for parasite growth and proliferation have not been explored. It was not surprising to find the chaperone HSP90 (named HSP83 in *Leishmania*) associated with TOR1 – many HSP90 clients are involved in signalling, and TOR and RPTOR1 have been identified as HSP90 interactors by others (http://www.picard.ch/Hsp90Int/index.php)^39^. Molecular chaperones coordinate with nutrient availability to regulate TORC1 assembly and signalling^40, 41, 42^. This allows TORC1 to rapidly respond to upstream signals and to link the stress response to TOR signalling for maintenance of protein homeostasis. In *Leishmania*, HSP83 is crucial for proliferation of both promastigotes and amastigotes and its inhibition causes growth arrest and promastigote to amastigote differentiation^43, 44, 45^. Furthermore, chaperones are expressed constitutively through the parasite’s various life-cycle stages but are phosphorylated stage-specifically with their phosphorylation correlating with specific complex formation that links to protein translation, growth and morphology^44, 46^. These findings suggest that they may be an important part of signalling pathways in response to nutrients and environmental cues including the TOR pathway.

RNAi of TOR1 and RAPTOR in the bloodstream form of *T. brucei* parasites results in cell cycle arrest, inhibition of protein synthesis and reduction in cell size, indicating that TORC1 is essential for trypanosome proliferation and growth^17^. Many *Leishmania spp* lack RNAi activity^47^ and we were unable to generate *RPTOR1* KO promastigotes through double homologous replacement and CRISPR-Cas9. We therefore made use of a conditional gene deletion system that relies on rapamycin-induced dimerization of the Cre subunits to study RPTOR1/TORC1 function^15^. In Opisthokonta, TOR signalling is assessed through phosphorylation of its substrates p70 S6 kinase and 4E-BP1 (restricted to Euteleostomi), which promote protein translation through ribosome biogenesis and translational initiation of capped mRNA, respectively^48, 49^. Orthologues of either of these proteins have not been identified in trypanosomatids nor have other TOR1 substrates been defined. This prevented us from directly assessing TORC1 signalling through phosphorylation of its targets, and we instead measured cellular activities associated with TORC1 signalling.

Our data showed that RPTOR1/TORC1 is an essential positive regulator of cell proliferation and growth of the insect-stage promastigotes. Loss of RPTOR1 induced proliferation arrest in the G1 phase of the cell cycle, inhibited protein synthesis and induced physiological changes characteristic of their differentiation to quiescent metacyclic promastigotes (Figure 7). RPTOR1 loss in other systems such as yeast (*S. cerevisiae*) and a range of different mammalian cell types results in a similar G1 cell cycle arrest and either inhibits cell cycle progression in proliferating cells or prevents cell cycle entry from quiescence^50, 51, 52^. We observed these effects in log-stage promastigotes, which differentiated to non-dividing metacyclic promastigotes and were unable to re-enter cell cycle to differentiate to retroleptomonads. Importantly, metacyclogenesis is not an automatic consequence of reduced proliferation. Serafim *et al* showed that depletion of adenosine or an adenosine receptor antagonist can inhibit proliferation and trigger metacyclogenesis in *Leishmania*, which can be reversed by addition of adenosine and other purines^24^. However, inhibiting the purine salvage pathway reduces proliferation but does not trigger metacyclogenesis. Conversely, supplementing parasites with a purine salvage pathway intermediate can induce proliferation while not interfering with metacyclogenesis^24^. Our work shows that deletion of *RPTOR1* produces metacyclic parasites even in nutrient rich medium, suggesting that purine or other nutrient sensing is upstream and provides a key activating signal to *Leishmania* TORC1, preventing metacyclogenesis under normal conditions. A reduction in TORC1 activity either through inducible deletion of *RPTOR1* or depletion of purines, inhibits proliferation and triggers the metacyclic differentiation program (Figure 7). It is also possible that *RPTOR1* deletion causes irreversible differentiation of parasites to metacyclic promastigotes that can no longer proliferate in response to nutrient availability through other pathways. On the contrary, overexpression of *RPTOR1* with potential overactivation of TOR1 results in a decrease in metacyclogenesis and more proliferation of cells when purines are available. Others have reported a similar correlation between RPTOR levels or TOR1 activation and cell proliferation. Overexpression of RPTOR in cancer cells increases TOR1 activation and cell proliferation while silencing of RPTOR has the opposite effect^53^. Chen *et al* showed that overactivation of mTOR in hematopoietic stem cells could drive cells from quiescence to increased proliferation resulting in premature ageing and reduced regenerative capacity^54^.

**Figure 7.**
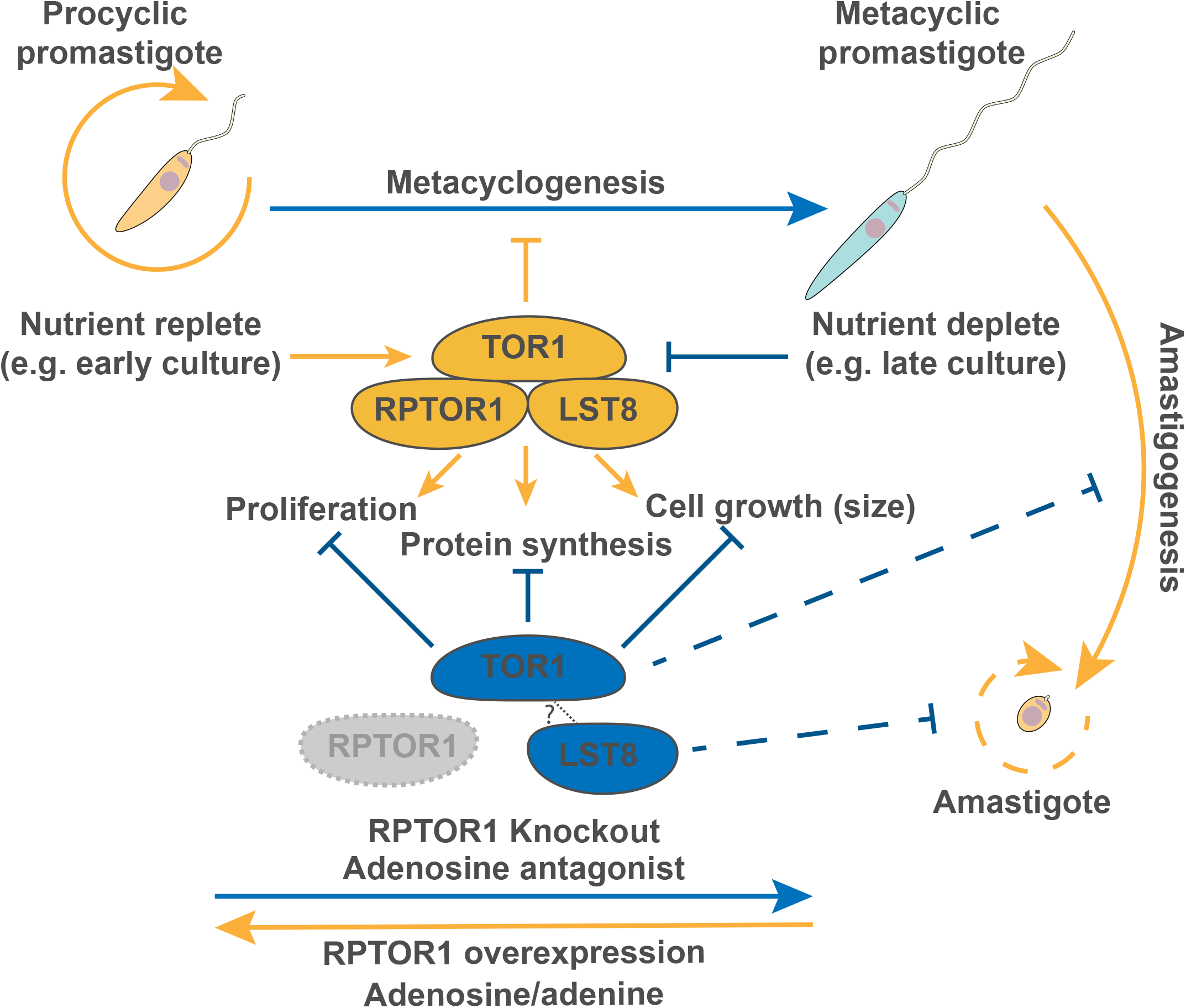
A proposed model of *Leishmania* proliferation or differentiation in response to nutrients and active TORC1. RPTOR1-dependent TORC1 is essential for the proliferation and growth of *Leishmania* procyclic promastigotes. Availability of nutrients (nutrient replete) activates TORC1 to promote parasite proliferation and growth while metacyclogenesis is inhibited. This can be enhanced through the supplementation of media with adenine (Figure 6G) or adenosine^24^, and/or through overexpression of RPTOR1 (Figure 6G). Conversely, when nutrients are depleted or TORC1 is inactivated through the deletion of RPTOR1, parasite proliferation and growth are inhibited (Figures 3 and 4) and metacyclogenesis occurs (Figure 6A-D). This results in the differentiation of procyclic promastigotes to metacyclic promastigotes that are unable to differentiate to proliferating retroleptomonads^27^ (Figure 6F) or develop into proliferating amastigotes in the murine host (Figure 6E).

RPTOR1 loss also prevented the expansion of amastigotes *in vivo* even though the metacyclics were capable of infecting macrophages *in vitro*. This provides evidence that TORC1 is important for the mammalian stage of the parasite and could potentially be targeted to inhibit growth and proliferation in their human or canine hosts. Further support for this comes from a recent study that investigated the Rag GTPases in the visceralizing species *Leishmania donovani*^32^. The Rag GTPases (RagA/C or RagB/D heterodimers) act upstream of TORC1 to sense amino acids, and the RagA/C complex is present in *Leishmania spp*. RagA was essential for promastigote growth while RagC was not essential for promastigotes but was required for parasite survival in mice. Unfortunately, little else is currently known about the TOR pathway in *Leishmania* but the bioinformatic analyses suggest distinct differences in sensing and signalling components that could be explored for selective chemotherapy.

RPTOR1 contains an N-terminal caspase-like domain and initial structure prediction analysis indicated conservation of the active-site cysteine-histidine (Cys-His) dyad suggesting that RPTOR1 may have peptidase activity^19^. However, structural data of the *Arabidopsis thaliana* (AtRAPTOR1) provided evidence that RAPTOR1 lacks the catalytic Cys-His dyad as the caspase cysteine is replaced by a serine while the adjacent cysteine faces into a hydrophobic core^20^. Our primary sequence alignment indicates that this may also be the case in kinetoplastids where the caspase cysteine is replaced by an asparagine (*Lm*, Asn^270^, Figure S4B). We used a complementation approach to show that the adjacent cysteine (Cys^269^) is not required to reverse the LmjRPTOR1 deletion phenotypes. RPTOR1 appears to be a pseudopeptidase, but the available structural data for human RPTOR1 show that the caspase-like domain is optimally positioned within the TORC1 active site cleft suggesting that it may be important for substrate recognition and recruitment^55^.

In summary, our data identify TORC1 as crucial for *Leishmania* growth and proliferation in response to nutrients, for both the insect-stage promastigotes and the mammalian-stage amastigotes. It also links the nutrient-sensing TORC1 pathway to differentiation to reveal TORC1 as a key mechanism by which *Leishmania* parasites inhibit differentiation from proliferating promastigotes to infective metacyclic promastigotes. This study sets a framework for further dissection of the TORC1 signalling cascade and its regulators.

## ACKNOWLEDGEMENTS

This work was supported by the Medical Research Council (MRC MR/K019384/1) and Wellcome Trust (200807) to J.C.M. We thank Karen Hogg, Graeme Park and Karen Hodgkinson for technical advice and support within the Bioscience Technology Facility, University of York; Nathaniel Jones, University of York, for generating the structural overlays with the LinfRPTOR1 Alphafold model and for critical review of the manuscript. The York Centre of Excellence in Mass Spectrometry was created thanks to a major capital investment through Science City York, supported by Yorkshire Forward with funds from the Northern Way Initiative, and subsequent support from EPSRC (EP/K039660/1; EP/M028127/1).

## AUTHOR CONTRIBUTIONS

Conceptualization, E.M., K.M., and J.C.M.; Methodology, E.M., V.G., E.V.C.A-F., K.M. and J.C.M.; Investigation, E.M., V.G., E.V.C.A-F., Y.R.N., J.S.G., E.B., and K.M.; Writing – Original Draft, E.M., V.G., and J.C.M.; Writing – Review & Editing, E.M., V.G., K.M. and J.C.M.; Funding Acquisition, J.C.M.; Resources, J.C.M.; Supervision, E.M. and J.C.M.

## MATERIALS AND METHODS

### Animals

Female BALB/c mice, 4–6 weeks old, purchased from Charles River Laboratories were used for animal studies. Mice were maintained under pathogen free conditions at the Biological Services Facility at the University of York (Heslington, York, UK). The studies were carried out in accordance with the Animal (Scientific Procedures) Act 1986 and under UK Home Office regulations using Project License 60/4442. Protocols and procedures were approved by the relevant ethics committees at the University of Glasgow and University of York.

### *Leishmania* parasites

*L. major* (MHOM/JL/80/Friedlin) and *L. mexicana* (MNYC/BZ/62/M379) were grown as promastigotes in HOMEM medium (modified Eagle’s medium, Invitrogen) supplemented with 10% (v/v) heat-inactivated fetal calf serum (Gibco) and 1% Penicillin/Streptomycin solution (Sigma-Aldrich) at 25◻°C. Transgenic parasite lines were cultured in the presence of appropriate antibiotics at the following concentrations: 50 μg ml^−1^ puromycin, 50 μg ml^−1^ hygromycin B, 10 μg ml^−1^ blasticidin and 25 μg ml^−1^ G418 (all from Invivogen).

## METHOD DETAILS

### Generation of cell lines

Inducible RPTOR1 null mutant (*RPTOR1^−/flox^*) lines were generated using a modified approach (Strategy 2 in^56^) of the DiCre inducible system^15^ (Figure S1B). Plasmids were constructed using Gateway cloning and transfected into log-stage *Leishmania major* promastigotes as previously described^15^. Briefly, a stable DiCre expressing cell line (DiCre) was generated by integrating genes for the two subunits of Cre-recombinase fused to FK506-binding protein (FKBP12) and the binding domain of the FKBP12-rapamycin associated protein into the 18S ribosomal RNA locus in *L. major* (Figure S2). This DiCre line was then used to generate an inducible RPTOR1 null mutant (*RPTOR1^−/flox^*) by replacing the first *RPTOR1* allele (LmjF.25.0610) with a LoxP flanked (floxed) C-terminal GFP fused RPTOR1 gene and replacing the second allele with a Hygromycin resistance cassette. Wildtype (WT) and C269A mutant RPTOR1 re-expression (addback) plasmids were generated by Gibson assembly and site-directed mutagenesis and transfected into an inducible *RPTOR1* null mutant (*RPTOR1^−/flox^*) line (cl2). For mass spectrometry analyses, *Raptor* (LmxM.25.0610), *TOR1* (LmxM.36.6320) and the control bait LmxM.29.3580 were N-terminally endogenously strep-tagged via CRISPR-Cas9 editing as previously described^16^. For co-immunoprecipitation, *TOR1* (LmxM.36.6320) was N-terminally endogenously myc-tagged in *L. mexicana* promastigotes using the same CRISPR-Cas9 editing approach. This latter line and the *L. mexicana* Cas9 T7 line were then transfected with the WT RPTOR1-HA addback plasmid to generate single or dual tagged lines. Oligonucleotides and plasmids used in this study are summarised in Table S2 and Table S3.

### Affinity purification of strep-RPTOR1 and strep-TOR1

Parasites expressing strep-RPTOR1, strep-TOR1 or the control bait LmxM.29.3580 were cultured to a density of 7.5×10^6^ parasites/ml. 7.5×10^8^ parasites per biological replicate were harvested by centrifugation for 10 min at 1200 x*g* and washed twice in PBS. Parasites were re-suspended at a density of 7.5×10^7^ parasites/ml in pre-warmed PBS. Dithiobis(succinimidyl propionate)(DSP, Thermo Fisher) cross-linker was added to a final concentration of 1 mM and cross-linking proceeded for 10 min at 26°C. DSP was quenched by adding Tris-HCl to 50 mM and incubating for 10 min shaking at RT. Parasites were centrifuged for 3 min at 1200 x*g* and frozen at −80°C. Each parasite pellet was lysed in 400 μL lysis buffer (1% IgePal-CA-630, 50 mM Tris pH 7.5, 250mM NaCl, 1mM EDTA, 0.1 mM PMSF, 1 μg ml^−1^ pepstatin A, 1 μM E64, 0.4mM 1-10 phenanthroline). Every 10 ml of lysis buffer was additionally supplemented with 200μl proteoloc protease inhibitor cocktail containing w/v 2.16% 4-(2-aminoethyl)benzenesulfonyl fluoride hydrochloride, 0.047% aprotinin, 0.156% bestatin, 0.049% E-64, 0.084% Leupeptin, 0.093% Pepstatin A (Abcam), 3 tablets complete protease inhibitor EDTA free (Roche) and 1 tablet PhosSTOP (Roche). Parasites were lysed by sonication with a microtip sonicator on ice for 3 rounds of 10 seconds each at an amplitude of 30. Insoluble material was pelleted by centrifugation at 10,000 x*g* for 10 min at 4°C. The cleared supernatant was added to 50 μL of Strep-Tactin XT resin and baits were affinity purified with end-over-end rotation for 2hrs at 4°C. Resin was washed 4x, using 300μL of ice cold lysis buffer for each wash, followed by 2x washes with 300μL ice cold PBS. Bait proteins were eluted in two rounds with 25 μL 50 mM biotin in 50 mM TEAB for 10 min each round. Proteins were precipitated with addition of 200 μL methanol then 50μL chloroform. After vortex mixing, proteins were pelleted by centrifugation at 18,000 x*g* for 1 hr at 4°C. The protein pellet was washed with ice cold methanol then resuspended in 200 μL 0.01% ProteaseMax in 50 mM TEAB, 10 mM TCEP, 10 mM IAA and 1 mM CaCl_2_. 200 ng of trypsin/lys-C (Promega) was added and proteins were digested overnight at 37°C. Digests were acidified by adding TFA to 0.5% and incubated at RT for 1 hr. After clarifying digests at 18,000 x*g* for 10 min, peptides were desalted using in house made C18 StageTips.

### Mass spectrometry data acquisition

Peptides were re-suspended in aqueous 0.1% trifluoroacetic acid (v/v) then loaded onto an mClass nanoflow UPLC system (Waters) equipped with a nanoEaze M/Z Symmetry 100 Å C_18_, 5 μm trap column (180 μm x 20 mm, Waters) and a PepMap, 2 μm, 100 Å, C_18_ EasyNano nanocapillary column (75 mm x 500 mm, Thermo). The trap wash solvent was aqueous 0.05% (v:v) trifluoroacetic acid and the trapping flow rate was 15 μL/min. The trap was washed for 5 min before switching flow to the capillary column. Separation used gradient elution of two solvents: solvent A, aqueous 0.1% (v:v) formic acid; solvent B, acetonitrile containing 0.1% (v:v) formic acid. The flow rate for the capillary column was 300 nL/min and the column temperature was 40°C. The linear multi-step gradient profile was: 3-10% B over 7 mins, 10-35% B over 30 mins, 35-99% B over 5 mins and then proceeded to wash with 99% solvent B for 4 min. The column was returned to initial conditions and re-equilibrated for 15 min before subsequent injections.

The nanoLC system was interfaced with an Orbitrap Fusion Tribrid mass spectrometer (Thermo) with an EasyNano ionisation source (Thermo). Positive ESI-MS and MS^2^ spectra were acquired using Xcalibur software (version 4.0, Thermo). Instrument source settings were: ion spray voltage, 2,100 V; sweep gas, 0 Arb; ion transfer tube temperature; 275°C. MS^1^ spectra were acquired in the Orbitrap with: 120,000 resolution, scan range: *m/z* 375-1,500; AGC target, 4e^5^; max fill time, 100 ms. Data dependent acquisition was performed in top speed mode using a 1 s cycle, selecting the most intense precursors with charge states >1. Easy-IC was used for internal calibration. Dynamic exclusion was performed for 50 s post precursor selection and a minimum threshold for fragmentation was set at 5e^3^. MS^2^ spectra were acquired in the linear ion trap with: scan rate, turbo; quadrupole isolation, 1.6 *m/z*; activation type, HCD; activation energy: 32%; AGC target, 5e^3^; first mass, 110 *m/z*; max fill time, 100 ms. Acquisitions were arranged by Xcalibur to inject ions for all available parallelizable time.

### Mass spectrometry data analysis

Peak lists in .raw format were imported into Progenesis QI (Version 2.2., Waters) and LC-MS runs aligned to the common sample pool. Precursor ion intensities were normalised against total intensity for each acquisition. A combined peak list was exported in .mgf format for database searching against the *L. mexicana* subset of the TriTrypDB database (8,250 sequences; 5,180,224 residues), appended with common proteomic contaminants (116 sequences; 38,371 residues). Mascot Daemon (version 2.6.0, Matrix Science) was used to submit the search to a locally-running copy of the Mascot program (Matrix Science Ltd., version 2.7.0). Search criteria specified: Enzyme, trypsin; Max missed cleavages, 1; Fixed modifications, Carbamidomethyl (C); Variable modifications, Oxidation (M), Phosphorylation (S,T); Peptide tolerance, 3 ppm; MS/MS tolerance, 0.5 Da; Instrument, ESI-TRAP. Peptide identifications were passed through the percolator algorithm to achieve a 1% false discovery rate assessed against a reverse database and individual matches filtered to require minimum expect score of 0.05. The Mascot .XML result file was imported into Progenesis QI and peptide identifications associated with precursor peak areas and matched between runs. Relative protein abundance was calculated using precursor ion areas from non-conflicting unique peptides. Accepted protein quantifications were set to require a minimum of two unique peptide sequences. Statistical testing was performed in Progenesis QI from ArcSinh normalised peptide abundances and ANOVA-derived p-values were converted to multiple test-corrected q-values using the Hochberg and Benjamini approach. Label free protein intensities were analysed with SAINTq^57^ to determine interacting proteins. Identified proteins in RPTOR1 and TOR1 affinity purifications were quantified relative to levels in the control bait (LmxM.29.3580) purification. Prey scores were filtered to achieve an overall false discovery rate of 10% in the final list of interactors.

### Functional annotation of genes identified by mass spectrometry

Proteins identified in the proteomic analyses were further annotated by homology-based methods and searches in TriTrypDB^58, 59^. A BLASTp^60^ search was performed against the NCBI Genbank non-redundant (NR) sequence database^61^ with an E-value threshold◻<◻= 1e-05.

### Co-immunoprecipitation

Co-immunoprecipitation was performed using TOR1-myc and RPTOR1-HA dual or single tagged log-stage promastigotes. 5×10^8^−10^9^ parasites were centrifuged at 1200 x*g* for 10 min, resuspended in 1 mL cold PBS and again centrifuged at 1200 x*g* for 5 min. Cells were then resuspended in PBS, 2mM DSP was added and cells were incubated for 10 min shaking at RT. This was followed by adding 50 mM Tris-Cl (final) pH-8 and incubating for 10 min shaking at RT. Cells were pelleted by centrifugation and resuspended in lysis buffer (20 mM Tris-Cl pH 8, 150 mM, NaCL, 0.5% NP-40, 2X mini complete Ultra (Roche), 0.5mM EDTA, 10 μM E-64. The cells were sonicated and lysed at 3x 30s 5W output on ice. The insoluble material was removed by centrifuging at 16000 x*g* for 10 min at 4°C and the supernatant was transferred into a fresh tube. 30 μL of HA Dynabeads (Thermo Fischer) were transferred to a 1.5 mL tube and washed 3x in lysis buffer using a magnetic rack for 2 min each wash. The cell extract was transferred on the beads and incubated for 1 h at 4°C under rotation. The beads were washed six times in 1 mL of wash buffer on a magnetic rack.

### Antibodies and western blotting

Polyclonal anti-RPTOR1 antibodies were raised in rabbits using an *L. major* recombinant RPTOR1 fragment consisting of the N-terminal 462 amino acids of the protein. The recombinant fragment was obtained by amplifying the relevant sequence from *L. major* genomic DNA, cloning the DNA fragment into pET28a(+) (Novagen) using oligonucleotides described in Table S2, and transforming into *Escherichia coli* BL21 DE3 (pLysS). The cells were then grown in LB medium containing 37 μg ml^−1^ chloramphenicol and 20 μg ml^−1^ kanamycin, until an A600 of 0.6 was reached and then induced with 1 mM isopropyl-β-d-thiogalactopyranoside overnight at 20 °C. The resulting 49 kDa protein was used for antibody generation in rabbits and the antibodies were affinity purified of from the rabbit sera.

For western blotting to confirm addback lines, 1×10^7^ rapamycin-induced cells were harvested and washed once in PBS by centrifugation at 1200 x*g*. Protein samples were prepared by lysing the pellet of parasites in laemmli buffer and boiling for 5 min; 5×10^6^ parasites were loaded per well. For co-immunoprecipitated samples, 50 μl of laemmli buffer was added to the beads after 6 washes, boiled for 5 min and 25 μl sample was loaded on an 8% NuPAGE Bis-Tris gel (Thermo scientific). Each gel was transferred to a PVDF membrane by wet transfer. The membrane was blocked in 5% milk for 1 hr, followed by incubation with rabbit anti-HA antibodies (1:3000, Bethyl), chicken anti-Myc antibodies (1:3000, 4A6 Merck), rabbit anti-RPTOR1 antibodies (1:200), or sheep anti-OPB antibodies (1:20000), 3 washes and incubation with HRP-conjugated secondary antibodies. The blots were developed using the SuperSignal West Pico substrate (Thermo Fisher).

### Induction of DiCre mediated *RPTOR1* deletion

In this DiCre system gene excision can be induced by addition of 100 nM to 1 μM rapamycin (Abcam), which dimerizes the Cre subunits resulting in an active diCre recombinase. Mid-log stage promastigotes were diluted to a density of 1 × 10^5^ cells mL^−1^ and induced for 3 days by daily addition of 100 nM rapamycin (Abcam) in DMSO, or DMSO alone (0.1%) in control cultures. After this initial induction, cells were counted and diluted in fresh supplemented HOMEM medium at a density of 1 × 10^5^ cells mL^−1^ with daily addition of 100 nM rapamycin until harvested for analysis.

### Diagnostic PCR to assess for presence of transgenes and *RPTOR1*

1-5 × 10^6^ cells were centrifuged at 1000 xg for 8 min, washed once in PBS and frozen at −20°C. DNA was extracted using the QIAGEN DNeasy Blood and Tissue Kit according to the manufacturer’s instructions for animal cells. All oligonucleotides used in this study are summarised in Table S2.

### Flow cytometry to analyse cell size, RPTOR1-GFP expression, viability and cell cycle

At the times indicated in figure legends cells were prepared for flow cytometry. Briefly, 5 × 10^6^ cells were centrifuged at 1000 x*g* for 8 min and washed twice in PBS with 5mM EDTA (PBS/EDTA). To assess cell size, viability, and RPTOR1-GFP expression, live cells were then resuspended in 1 mL of PBS/EDTA with 1 μg mL^−1^ of propidium iodide (PI) to allow assessment or exclusion of dead cells. For cell cycle analysis cells were fixed in 70% methanol in PBS/EDTA at 4°C for 1 hr or overnight and washed twice in PBS/EDTA by centrifugation at 1000 xg for 5 min. Cells were resuspended in 1 mL of PBS/EDTA with 10 μg mL^−1^ of PI and 10 μg mL^−1^ of RNAseA, and incubated at 37°C for 45 min. Fixed or live cells were analysed on a Beckman Coulter, CyAn ADP and data were analysed on FlowJo software (Tree Star Inc.).

### Cell proliferation and clonogenic survival assay

To assess proliferation, cells were counted daily using a hemocytometer or Z1 Beckman Coulter counter before and after rapamycin induction. For the clonogenic survival assay, cells were induced for 72 hr with rapamycin or DMSO, counted and diluted to 1.6 parasites mL^−1^ in HOMEM with 20% heat-inactivated FCS and 100 nM rapamycin or DMSO. The cells were plated out in two to four 96-well flat bottom plates by adding 200 μL cells per well. Plates were sealed and incubated at 25°C for 3-4 weeks. Surviving clones were counted by visual inspection of each well using a light microscope; any well containing live parasites was counted as a surviving clone. The percentage of surviving clones of the total cells plated is shown in graphs.

### Protein synthesis

Protein synthesis was assessed using a Click-It AHA Alexa Fluor 488 Protein Synthesis HCS Assay (Thermo Fisher, Cat# C10289). Click-iT reagents were prepared according to the manufacturer’s instructions. Cells were centrifuged at 1000 xg for 8 min and washed once in methionine-free RPMI medium supplemented with 10% of 3.5 kDa-dialysed heat inactivated FBS, 1 M HEPES (pH 7.4), 5 mM Adenine (Sigma A9126 Adenine Hemisulfate salt), 0.25% (2.5 mg ml^−1^) Hemin in 50% triethanolamine (Sigma H5533) or in 50 mM NaOH, 200 mM L-glutamine (Gibco), Pen/strep (Gibco) and 0.3 mg mL^−1^ Biopterin (Methionine-free medium, MFM). Cells were then resuspended in MFM containing 50 μM Click-iT® AHA working solution, transferred to 1.5 mL tubes (3 × 10^6^ cells per tube) in triplicate and incubated for 1-2 hours. After incubation, cells were centrifuged at 1000 xg for 5 min, washed once in PBS and fixed with 1% formaldehyde in PBS for 15 min at room temperature. Cells were then washed twice in 3% BSA in PBS and permeabilized using 0.1% Triton-X100 for 10 min at room temperature. Permeabilized cells were washed twice in 3% BSA in PBS, resuspended in 100 μL Click-iT reaction cocktail and incubated for 30 min at room temperature, protected from light. Cells were then washed once in 3% BSA in PBS and once in PBS alone by centrifugation at 1000 x*g* for 5 min, and finally transferred to a black 96-well flat-bottomed plate in 100 μL PBS. Alexa-Fluor 488 fluorescence was measured using a Clariostar microplate reader (BMG).

### Sequence alignment and topology diagrams

Sequences of caspase 7 and RPTOR1 from human and *A. thaliana* were obtained from UniProtKB (P55210, Q8N122 and Q93YQ1, respectively). *L. major* and *T. brucei* RPTOR1 sequences were obtained from TriTrypDB^58, 59^. The alignment of primary protein sequences was done using Clustal Omega 3^62^ and ALINE^63^; secondary structure alignment was done using PDBeFold from the structures of human Caspase_7 (PDB code 1F1J) and *A. thaliana* RAPTOR1 (PDB code 5WBI). STRIDE^64^ was used to assign secondary structure in the topology diagrams and the diagrams were produced with TOPDRAW^65^. The Alphafold model of LinfRPTOR1 was last updated in AlphaFold DB version 2022-06-01 and created with the AlphaFold Monomer v2.0 pipeline. AtRAPTOR1 and LinfRPTOR1 were superposed in UCSF Chimera (V1.14) using the MatchMaker tool (Needleman-Wunsch Algorithm, BLOSUM-62 Matrix and default parameters). The Match->Align tool was used to generate amino acid alignments from the structural superposition.

### Assessment of metacyclogenesis and retroleptonomad growth

PNA^−^ cells were isolated from cultures by agglutination of promastigotes with 50 μg ml^−1^ peanut lectin as previously described^23^. SHERP expression was measured using flow cytometry after staining with affinity-purified rabbit anti-SHERP^66^ and goat anti-rabbit AF647 antibodies. Briefly, cells were washed in PBS by centrifugation at 1000 x*g* for 3 min and fixed in 1% paraformaldehyde in PBS at 4°C overnight. Cells were washed again in PBS and permeabilized in 0.1% Triton X-100 for 5 min. After another three washes in PBS, cells were blocked for 30 min in PBS containing 10% FCS and 5% goat serum followed by incubation for 1 hr with rabbit anti-SHERP antibody (1/100 dilution) in blocking buffer, both on ice. Cells were washed three times in PBS and stained with goat anti-rabbit AF647 in PBS for 30 min on ice. Stained cells were washed twice in PBS and resuspended in PBS with 5 mM EDTA before analysis on a Beckman Coulter, CyAn ADP. Data were analysed on FlowJo software (Tree Star Inc.). To assess growth of retroleptomonads, promastigotes were purified twice using PNA agglutination as described above and resuspended in HOMEM containing 20% FCS followed by daily counting of live cells.

### Macrophage infections

Macrophages were generated from isolated bone marrow cells. Cell suspensions were obtained from femurs and tibia of BALB/c mice and cultured for six days in DMEM supplemented with 20% FCS, 2 mM L-glutamine, 100 U ml^−1^ penicillin, 100 μg ml^−1^ streptomycin and 30% L-929 conditioned medium (BMM medium). Fresh medium was added on day 3 and replaced with BMM medium without L-929 conditioned medium on day 6. On day 7 or 8, cells were harvested and replated in RPMI supplemented with 10% FCS in 16-well Nunc Lab-Tek chamber slides (Thermo Fisher) for use in infection assays. PNA^−^ promastigotes were added to bone-marrow macrophages at a 1:1 ratio and washed off after 24 hours using warmed RPMI supplemented with 10% FCS, 2 mM L-glutamine, 100 U ml^−1^ penicillin, 100 μg ml^−1^ streptomycin. On day 1, 3 and 5 after infection slides were washed with PBS, fixed with 3% paraformaldehyde, permeabilized with 100% methanol and stained with DAPI to assess infection using fluorescence microscopy.

### Lesion development in mice

Female BALB/c mice (4–6 weeks, Charles River Laboratories) were infected intradermally in the ear with 1×10^5^ PNA^−^ metacyclic promastigotes in 10 μl PBS. Lesion development was monitored weekly by using the Schuster scoring system^67^.

### Scanning electron microscopy

Rapamycin or DMSO treated cells were fixed in 4% formaldehyde and 2.5% glutaraldehyde in 0.1M phosphate buffer, pH 7.3 for 30 min. Cells were washed twice in 0.1M phosphate buffer for 10 min, adhered to poly-L-lysine-coated coverslips and post-fixed in 1% osmium tetroxide for 45 min on ice. Thereafter, they were washed twice in 0.1 M phosphate buffer for 10 min and dehydrated in a graded series of ethanol concentrations (25-100%) for 15 min each. The final 100% ethanol was replaced with two changes of hexamethyldisilazane (HMDS), and cells were left to air-dry in a desiccator overnight. Samples were affixed to SEM stubs, sputter coated with 20nm of gold-palladium on Polaron SC7640 sputter coater and then imaged using a JEOL JSM 6490LV scanning electron microscope operating at 8kV accelerating voltage. Images were analysed on ImageJ software (Fiji plugin).

### Assessment of adenine response

DiCre, *RPTOR1^−/flox^* Cl2 and *RPTOR1* complementation lines were cultured for five days in Grace’s insect medium (Sigma-Aldrich) with 10% heat-inactivated FCS (Gibco) and 1% Penicillin/Streptomycin solution (Sigma-Aldrich) with daily addition of 100 nM rapamycin (Abcam) in 0.1% DMSO to induce RPTOR1 excision, or 0.1 % DMSO alone in control cultures. After the first three days of culture cells were diluted to 1 × 10^5^ cells ml^−1^ in fresh media with rapamycin or DMSO and cultured for the remaining two days. After induction, cells were counted, diluted to a density of 2 × 10^6^ cells ml^−1^ and cultured in fresh Grace’s medium also supplemented with 500 μM adenine or DMSO as a control. After three days of culture, the cells were counted using a Z1 Beckman Coulter counter.

### Statistical analyses

All statistical analyses were performed with Prism (GraphPad Software, La Jolla, CA, USA) using the test specified in the figure legends. Statistically significant differences (P < 0.05) are annotated on the graphs as described in the figure legends.

## SUPPLEMENTAL INFORMATION

Document S1 – Figures S1-S6.

Table S1. Proteins identified in pull downs and associated SAINTq scores. Related to Figure 1.

Table S2. Oligonucleotides used in the study.

Table S3. Plasmids used in the study.

