## Supplementary material for "TORC1 is an essential regulator of nutrient-dependent differentiation in *Leishmania*": Document S1 - Figures S1-S6

### Figure S1

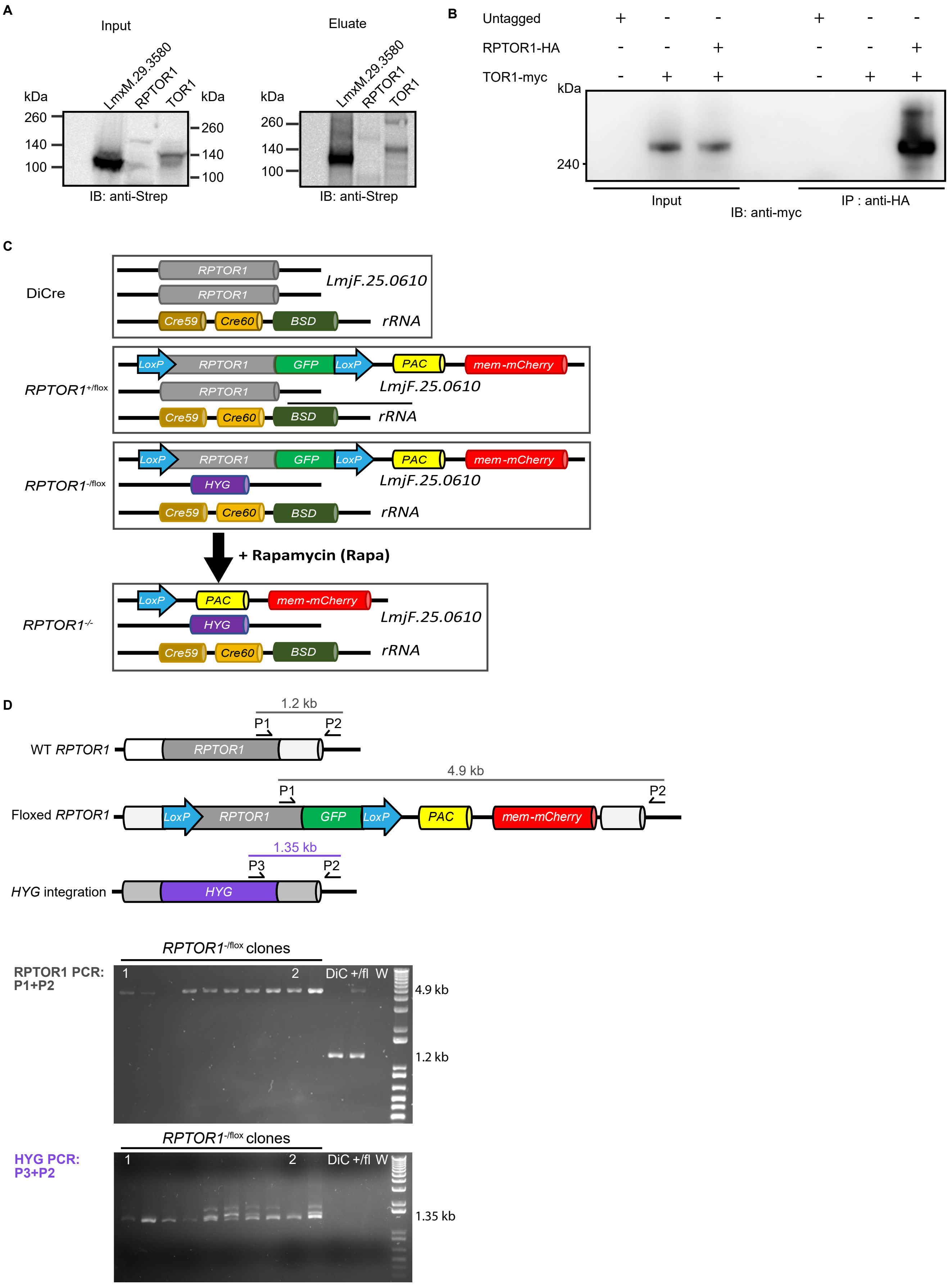

A
Input
Eluate
kDa
LmxM.29.3580
RPTOR1
TOR1
260
140
100
IB: anti-Strep
B
Untagged
RPTOR1-HA
TOR1-myc
240
Input
IP : anti-HA
IB: anti-myc
C
DiCre
RPTOR1
LmjF.25.0610
Cre59
Cre60
BSD
rRNA
LoxP
GFP
PAC
mem-mCherry
RPTOR1+/flox
RPTOR1-/flox
HYG
+ Rapamycin (Rapa)
RPTOR1-/-
D
1.2 kb
P1
P2
WT RPTOR1
4.9 kb
Floxed RPTOR1
1.35 kb
P3
HYG integration
RPTOR1-/flox clones
RPTOR1 PCR: P1+P2
DiC +/fl W
HYG PCR: P3+P2

**Figure S1. RPTOR1 immunoprecipitation and knockout strategy. Related to Figures 1 and 2**

**(A)** Affinity purification of endogenously Twin-Strep-tagged RPTOR1, TOR1 and control bait LmxM.29.3580. A sample of lysate from the parental line (T7) or Strep-tagged lines was taken prior to affinity purification (input) and analysed by western blot. Bait proteins were eluted from streptactin beads with biotin and a sample taken for analysis by western blot (eluate), with the remainder analysed by mass spectrometry. Predicted sizes are: LmxM.29.3580 85kDa, RPTOR1 161kDa, TOR1 291kDa. **(B)** Lysates of *L. mexicana* expressing untagged RPTOR1 and TOR1, HA-tagged RPTOR1 and/or myc-tagged TOR1 were incubated with anti-HA-conjugated magnetic beads and analysed by western blot using anti-myc antibodies. **(C)** Schematic of *RPTOR1* knockout strategy. The background cell line, DiCre, was generated by integrating the diCre expression cassette into the ribosomal RNA locus for constitutive expression of FKBP-Cre59 and FRB-Cre60 in *L. major* Friedlin. The inducible *RPTOR1* knockout line (*RPTOR1*$^{-/flox}$) was generated by replacing the 1st *RPTOR1* allele with a LoxP flanked (floxed) C-terminal GFP-tagged version of *RPTOR1* followed by replacement of the 2nd allele with a hygromycin resistance cassette. The floxed *RPTOR1* gene can be excised by Cre-recombinase following rapamycin induced dimerization to generate a *RPTOR1*$^{-/-}$ line. **(D)** Diagnostic PCRs of gDNA from generated cell lines confirm integration of hygromycin resistance and floxed *RPTOR1* cassettes. Primer binding site and size of PCR products are shown in the diagram. Ten clones of *RPTOR1*$^{-/flox}$ are shown after PCR and agarose gel electrophoresis (bottom part); lanes with clones 1 and 2 that are described in this study are indicated on the gel image. DiC, DiCre; +/fl, *RPTOR1*$^{+/flox}$; W, water control.

**Figure S2**

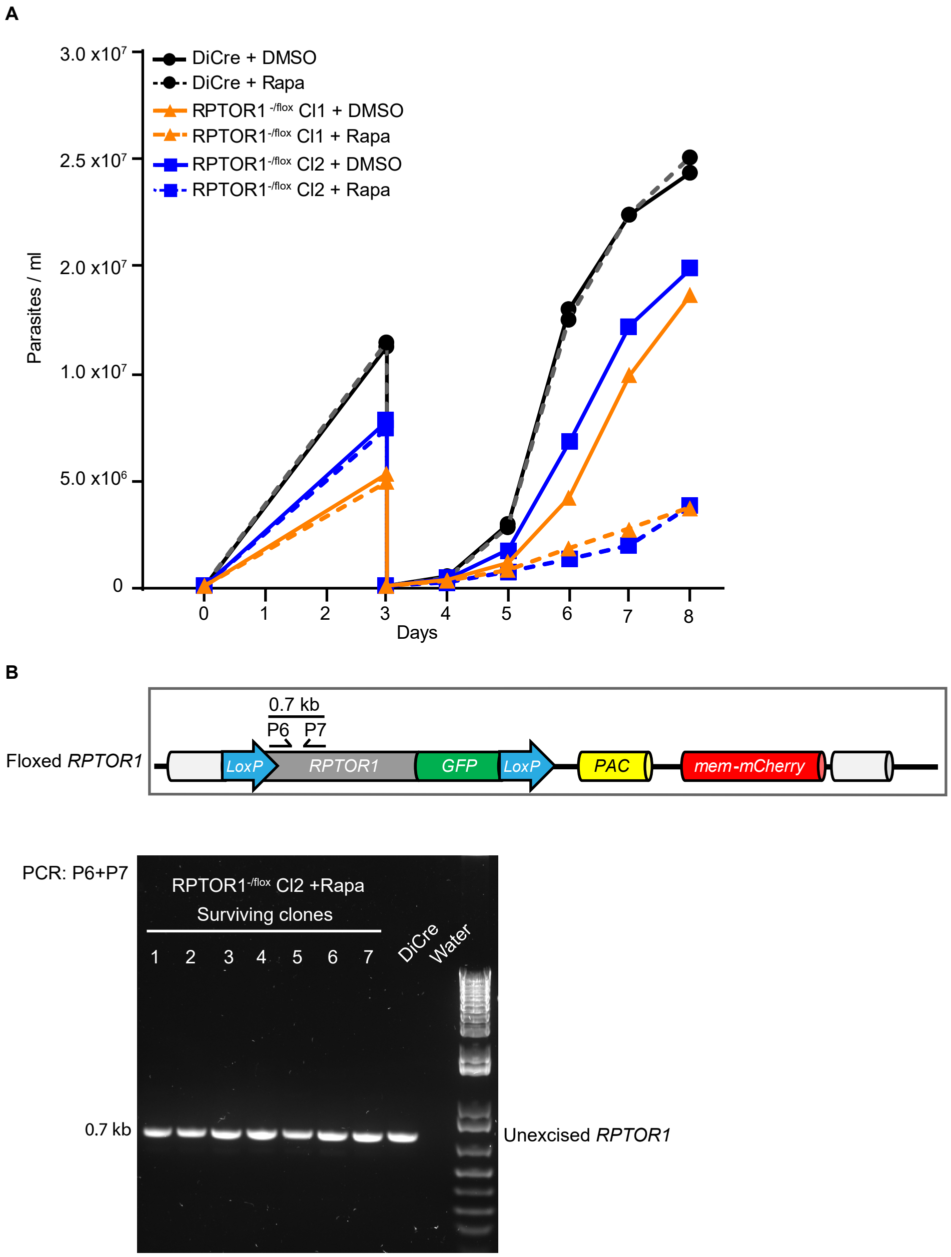


**Figure S2. RPTOR1 is essential for cell proliferation and long-term survival. Related to Figure 3.**

**(A)** Cell densities of uninduced (+DMSO, solid line) and rapamycin-induced (+Rapa, dashed line) cells. Log-stage promastigotes of DiCre (black) and RPTOR1-/flox lines, Cl1 (orange) and Cl2 (blue), were set up at $1x10^5$ cells mL-1 (day 0) and treated for three days with daily addition of DMSO or 100 nM rapamycin; cells were then counted and diluted to $1x10^5$ cells mL-1 (day 3) followed by culturing and daily counting for five days. A representative dataset of three to four similar experiments is shown. **(B)** PCR analysis of genomic DNA from surviving clones (clones 1-7) of rapamycin-induced $RPTOR1^{-/flox}$ Cl2 and DiCre cells. Schematic (upper panel) shows the RPTOR1 locus with floxed RPTOR1 cassette with primer binding sites and the predicted length of the PCR amplicon. Agarose gel (lowel panel) indicates the presence of the RPTOR1 CDS fragment from the unexcised floxed RPTOR1 cassette in the seven clones from 2 independent clonogenic assays.

Figure S3

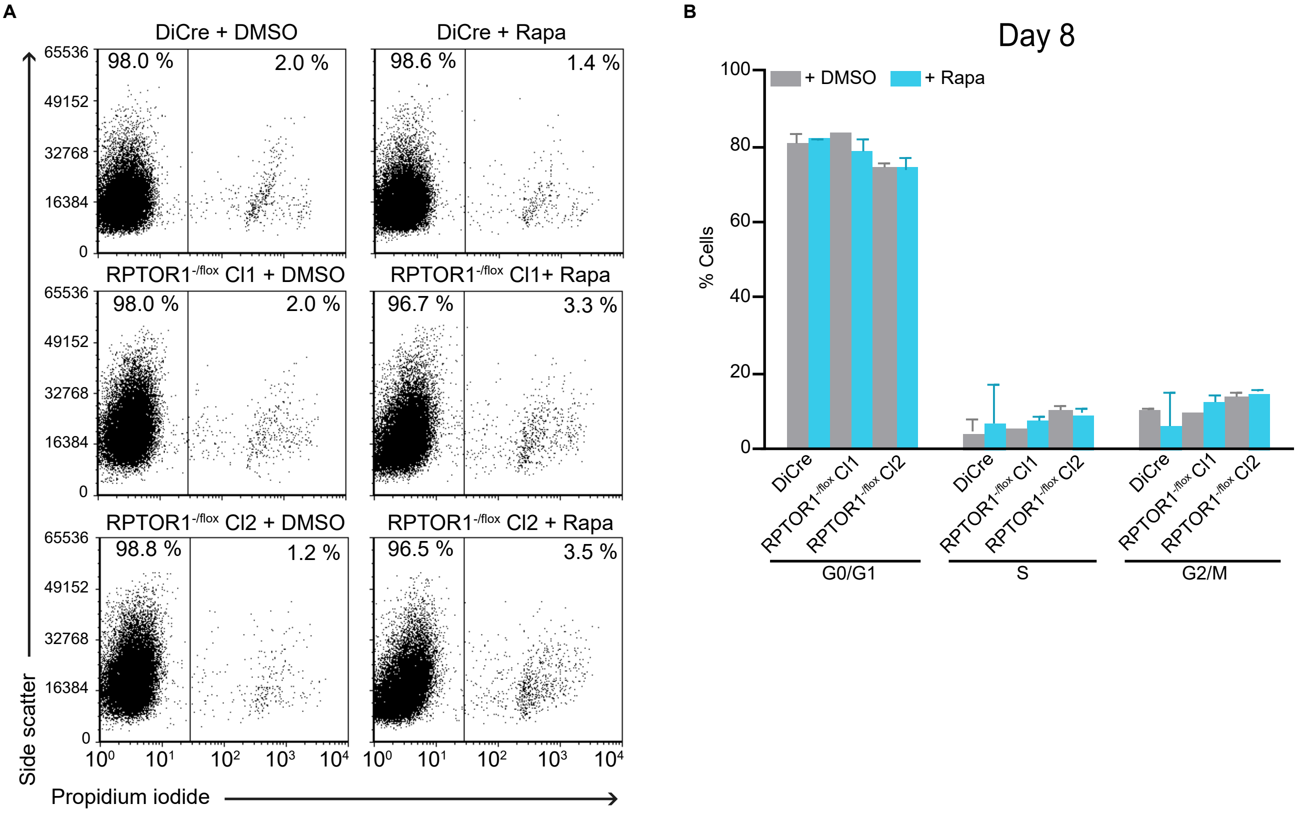


**Figure S3. Flow cytometry to assess viability and cell cycle defects. Related to Figure 3.** **(A)** Cell viability was measured by flow cytometry of propidium iodide-stained cells after five days of induction. Representative dot plots of side scatter versus propidium iodide fluorescence are shown for uninduced (+DMSO) or rapamycin-induced (+Rapa) cells. Numbers indicate the percentages of cells within the gate with live cells shown in the propidium iodide negative (left) gate in each plot. **(B)** Cell cycle analysis of fixed propidium iodide cells after eight days of induction. In both (A) and (B) cells were diluted after the first three days of induction and cultured for the remaining time with addition of more DMSO or rapamycin.

#### Figure S4

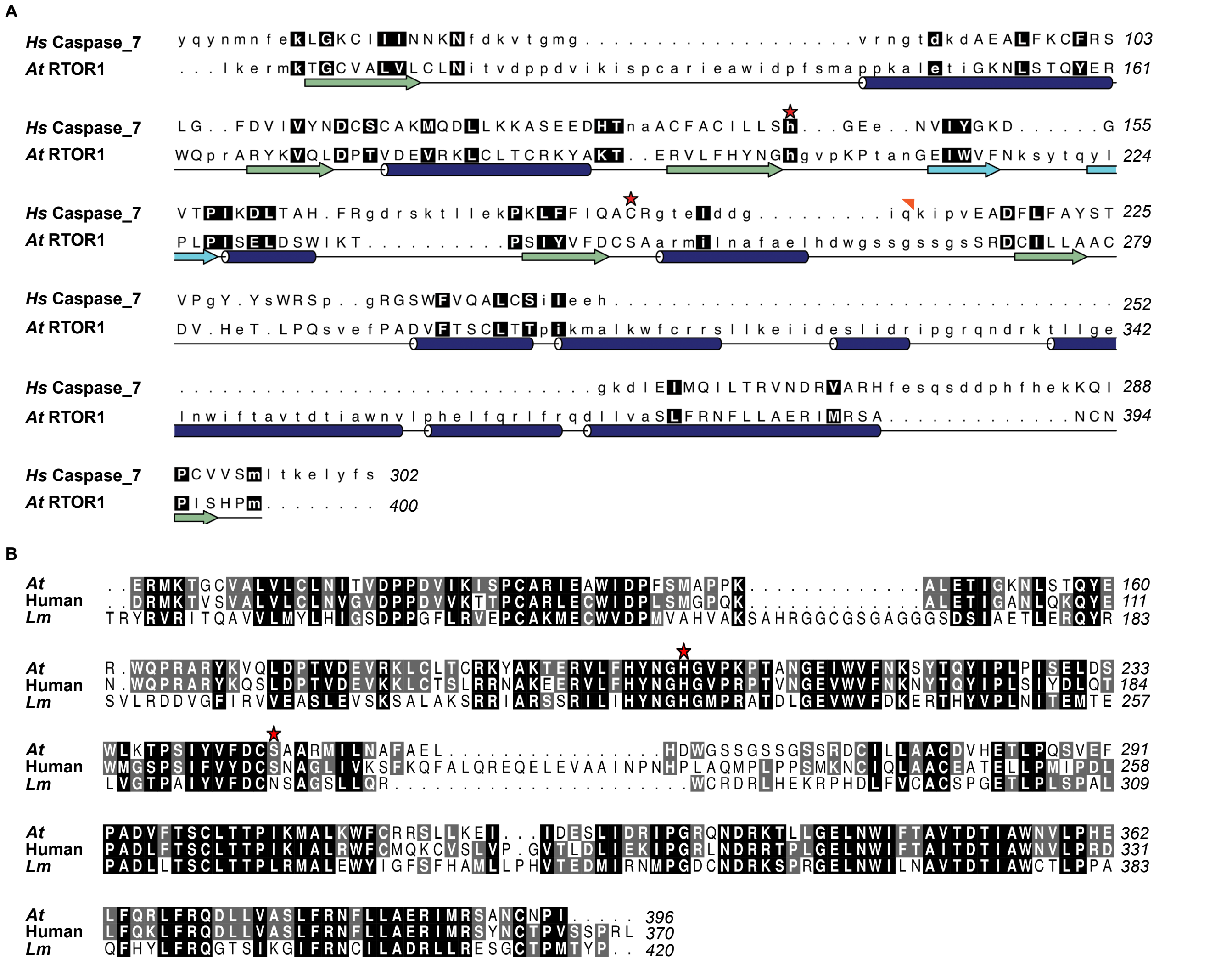

**Figure S4. Primary and secondary sequences alignments with RPTOR1. Related to Figure 5.**

**(A)** Sequence alignment based on structure and secondary structure elements of human caspase-7 and the RNC from *A. thaliana* RAPTOR1. β-strands are coloured green; the histidine and cysteine residues of the dyad is indicated for caspase-7 by red stars. The position where residues are missing from the caspase 7 structure (cleaved loop) is indicated by an orange triangle. Regions where the structures overlay well are shown in upper case while those whose backbone deviates more or don't align are in lower case. Numbers at the end of each line indicate the position of the last residues in that line in the UniProt sequences. **(B)** Primary amino acid sequence alignment of *A. thaliana* RAPTOR1, human RPTOR and *L. major* RPTOR1. White letters on black background indicate identical residues and white letters on a grey background indicate similar residues; the residues corresponding to the histidine cysteine dyad in caspase-7 shown in (A) are indicated by red stars.

**Figure S5**

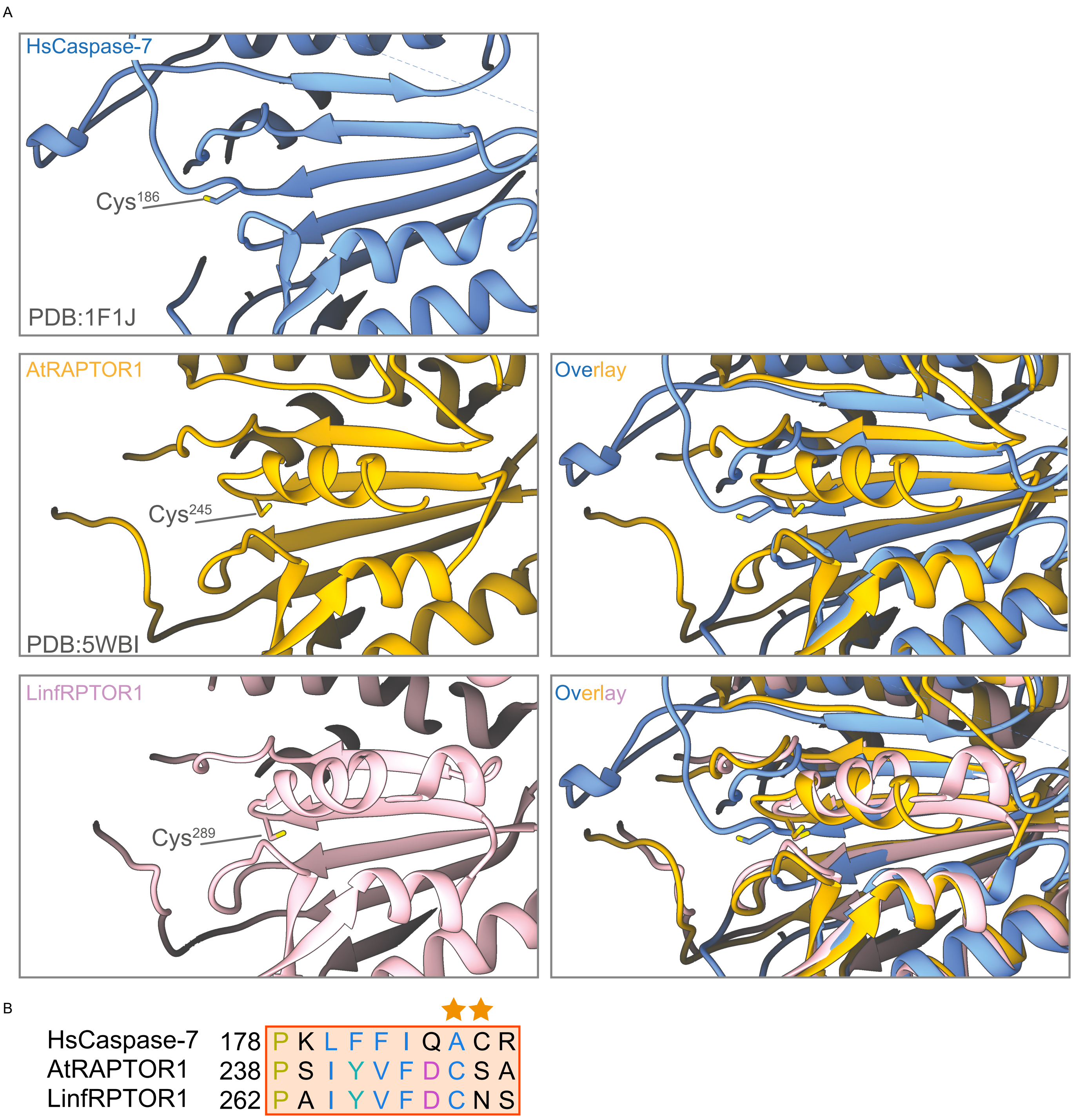


**Figure S5. Secondary sequences alignments using Alphafold model of RPTOR1. Related to Figure 5.**

**(A)** The X-ray crystal structures of HsCaspase-7 (PDB:1F1J), AtRAPTOR1 (PDB:5WBI), and the Alphafold model of LinfRPTOR1 (LINF_250011400) are shown individually on the left hand panels. The active site cysteine residues are denoted by the labels. AtRAPTOR1 and LinfRPTOR1 were superposed in UCSF Chimera using the MatchMaker tool (right hand panels). **(B)** The Match->Align tool was used to generate amino acid alignments from the structural superposition. Residues are coloured using the ClustalX scheme. The active site cysteines are annotated by stars above the sequence.

### Figure S6

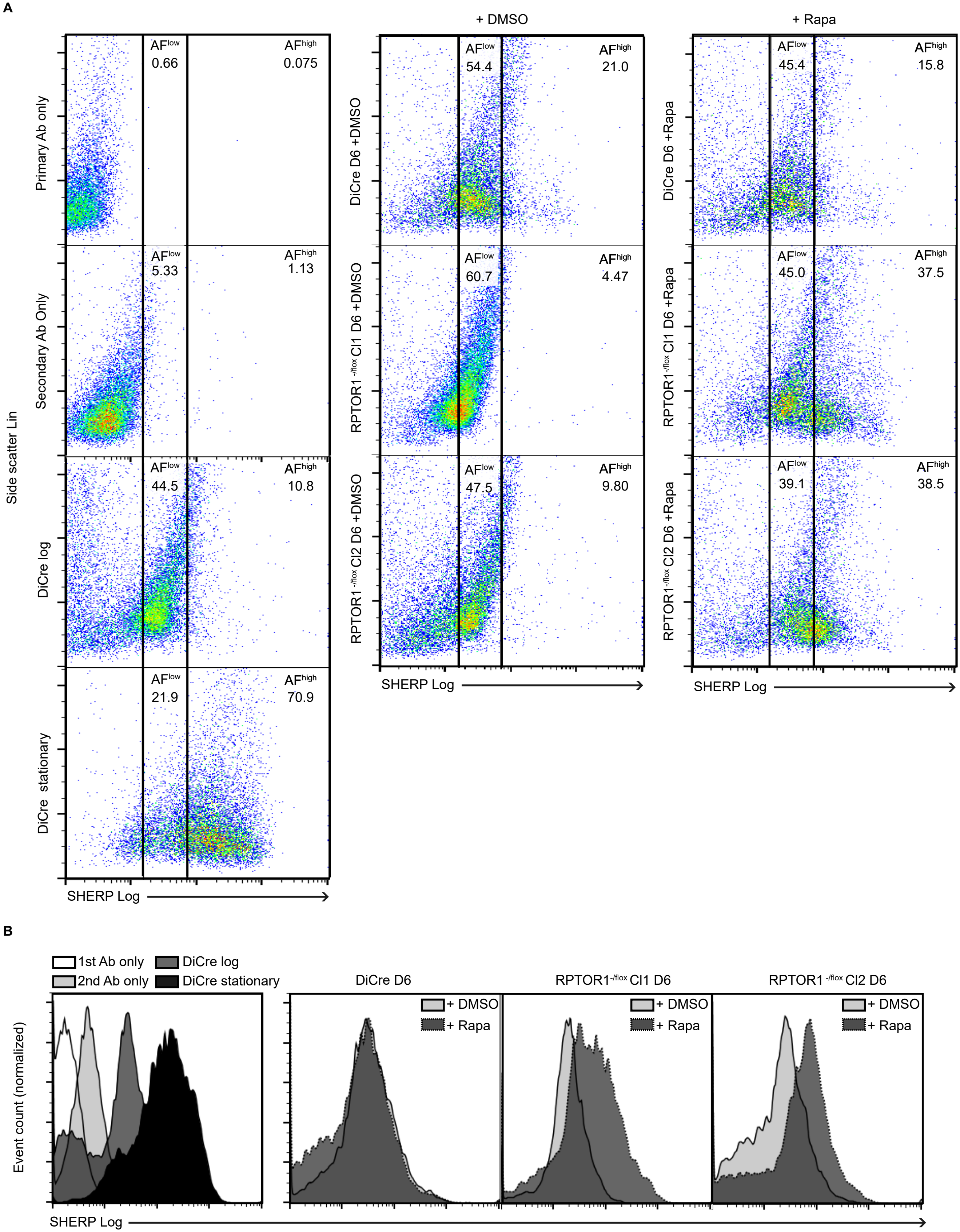
A
+ DMSO
+ Rapa
Primary Ab only
Secondary Ab Only
DiCre log
DiCre stationary
Side scatter Lin
AFlow 0.66
AFhigh 0.075
AFlow 5.33
AFhigh 1.13
AFlow 44.5
AFhigh 10.8
AFlow 21.9
AFhigh 70.9
DiCre D6 +DMSO
AFlow 54.4
AFhigh 21.0
RPTOR1 -/flox Cl1 D6 +DMSO
AFlow 60.7
AFhigh 4.47
RPTOR1 -/flox Cl2 D6 +DMSO
AFlow 47.5
AFhigh 9.80
DiCre D6 +Rapa
AFlow 45.4
AFhigh 15.8
RPTOR1-/flox Cl1 D6 +Rapa
AFlow 45.0
AFhigh 37.5
RPTOR1-/flox Cl2 D6 +Rapa
AFlow 39.1
AFhigh 38.5
SHERP Log
B
1st Ab only
2nd Ab only
DiCre log
DiCre stationary
DiCre D6
RPTOR1-/flox Cl1 D6
RPTOR1 -/flox Cl2 D6
+ DMSO
+ Rapa
Event count (normalized)
SHERP Log

**Figure S6. Flow cytometry analysis of SHERP expression. Related to Figure 6.**
SHERP expression was measured by flow cytometry after staining with anti-SHERP and Alexa Fluor 647 (AF647)-conjugated secondary antibodies. Log-stage promastigotes were treated for three days with DMSO or rapamycin (+Rapa), diluted and cultured for three more days (D6) with daily DMSO and rapamycin treatment. **(A)** Representative dot plots of side scatter versus AF647 fluorescence signal (SHERP staining) are shown. Controls (left panel) include DiCre cells stained with primary (1st Ab) or secondary antibody (2nd Ab) only and SHERP-stained DiCre early-log (D6, log) or stationary-phase cells. **(B)** Histograms of AF647 fluorescence (SHERP staining) in control (left panel) and DMSO or rapamycin treated cells.
